# Picky with peakpicking: assessing chromatographic peak quality with simple metrics in metabolomics

**DOI:** 10.1101/2023.07.28.551024

**Authors:** William Kumler, Bryna J. Hazelton, Anitra E. Ingalls

**Affiliations:** University of Washington

## Abstract

**Background:** Chromatographic peakpicking continues to represent a significant bottleneck in automated LC-MS workflows. Uncontrolled false discovery rates and the lack of manually-calibrated quality metrics require researchers to visually evaluate individual peaks, requiring large amounts of time and breaking replicability. This problem is exacerbated in noisy environmental datasets and for novel separation methods such as hydrophilic interaction columns in metabolomics, creating a demand for a simple, intuitive, and robust metric of peak quality.

**Results:** Here, we manually labeled four HILIC oceanographic particulate metabolite datasets to assess the performance of individual peak quality metrics. We used these datasets to construct a predictive model calibrated to the likelihood that visual inspection by an MS expert would include a given mass feature in the downstream analysis. We implemented two novel peak quality metrics, a custom signal-to-noise metric and a test of similarity to a bell curve, both calculated from the raw data in the extracted ion chromatogram and found that these outperformed existing measurements of peak quality. A simple logistic regression model built on two metrics reduced the fraction of false positives in the analysis from 70-80% down to 1-5% and showed minimal overfitting when applied to novel datasets. We then explored the implications of this quality thresholding on the conclusions obtained by the downstream analysis and found that while only 10% of the variance in the dataset could be explained by depth in the default output from the peakpicker, approximately 40% of the variance was explained when restricted to high-quality peaks alone.

**Conclusions:** We conclude that the poor performance of peakpicking algorithms significantly reduces the power of both univariate and multivariate statistical analyses to detect environmental differences. We demonstrate that simple models built on intuitive metrics and derived from the raw data are more robust and can outperform more complex models when applied to new data. Finally, we show that in properly curated datasets, depth is a major driver of variability in the marine microbial metabolome and identify several interesting metabolite trends for future investigation.

## Background

Liquid chromatography-mass spectrometry (LC-MS) is a powerful tool for exploring the molecular composition of biological samples. Its rapid sample processing (typically <1 hr run time), low limits of detection (pM-nM range), and ability to characterize novel molecules via fragmentation fingerprints make it a common workhorse for metabolomic research. In the past two decades, data-driven methods have established workflows for untargeted metabolomics but the imperfect performance of the core peakpicking algorithms continue to require manual oversight and curation. This problem has been exacerbated by the increased use of non-traditional chromatography such as hydrophilic interaction which tends to produce noisier peaks (Bajad et al. 2006; Myers et al. 2017b; Gika et al. 2019).

Noisy data and imperfect detection algorithms introduce a tradeoff between false positives (where contamination, background instrument or chemical noise is misclassified as biological signal) and false negatives (where real signals are undetected). Existing algorithms tend to favor the inclusion of false positives because downstream analyses can always remove erroneous mass features, but false negatives cannot be later recovered (Pirttilä et al. 2022; Gloaguen, Kirwan, and Beule 2022). However, this approach requires more time from the researcher as they manually evaluate a potentially enormous number of mass features (MFs), a task that scales combinatorially with the number of samples and compounds measured (Myers et al. 2017a). Instead of minimizing false negatives, we believe that emphasis should be placed on allowing the experimenter to set a threshold for the proportion of false positives (the false discovery rate or FDR) and accept that this will inherently add to the number of MFs already lost in the data collection process.

Existing peak-detection softwares do not provide a clear way to exclude false positives in an a consistent and unbiased way. Typical outputs consistent across the different implementations consist of the *m/z* ratio, retention time, and area for each mass feature, with some additional useful information occasionally provided such as the peak’s signal-to-noise ratio or degree of skew (Pirttilä et al. 2022). None of these parameters answer the critical question about the likelihood that a given feature corresponds to a molecule present in the original sample. This parameter is crucial for downstream analysis because it represents the base rate for error propagation and acceptable thresholds should vary widely by the particular project’s goals. In an exploratory analysis, any mass feature more than 50% likely to be real is perhaps worth considering, while in a confirmatory study this threshold may need to be above 99% likely to be real. Despite significant effort invested in improving the peakpicking algorithms, very little has been done to quantify the accuracy and precision of their outputs across the wide variety of datasets to which they are applied.

A single parameter of MF quality also facilitates downstream analyses in multiple ways. This metric would improve statistical power by reducing the number of effective hypotheses tested and allow researchers to focus effort on features least likely to be noise. Additionally, this parameter could be optimized to improve peakpicking and chromatographic settings independently of the software used and minimize inter-lab variability when scripted to provide consistent, reproducible results independent of the particular expert reviewing its performance. Constructing such a single comprehensive metric calibrated to likelihood is also more effective than multiple independent thresholds because it has meaningful units, does not require estimating the relative power of individual metrics, and allows a good MF to compensate for weak performance in one area with strong performance in other metrics, e.g. as implemented in Pirttilä et al. (2022) and Kantz et al. (2019).

An area particularly ripe for improved tools for metabolomic data analysis is that of the open ocean (Kido Soule et al. 2015). Low compound and high salt concentrations make metabolomics analyses difficult to study in this area but its vast size and the direct effect of its microbial communities on the Earth’s biogeochemistry make it critical that we understand the transformation of energy and nutrients on a molecular scale (Boysen et al. 2018). Metabolites are the currency of chemical exchange both intra- and inter-cellularly, serving as building blocks of larger molecules, regulators of osmotic balance and storage of nutrients, as well as important chemical signals on their own. These small molecules serve both as signposts for the complex biological landscape in this highly dynamic region and give a sense of not only who is present but also what ecological roles they’re serving and the niches they fill (Kido Soule et al. 2015; Boysen et al. 2021; Heal et al. 2021).

In this paper, we use open ocean marine metabolite LC-MS samples to develop and test a variety of chromatographic peak metrics. We construct and validate multiple predictive models of MF quality based on metrics both common in the literature and custom implementations we’ve found useful in our own analysis. This allows us to connect the physical, chemical, and biological measurements taken regularly around the globe to a molecular-scale perspective of particulate organic matter in the ocean by linking the chemical currencies that fuel the planet to the environments in which they’re found.

## Results

### Dataset characterization

An average of 3,300 mass features (MFs) were reported by XCMS across the 4 datasets, with the fewest (1,495) in the Falkor data and the most in the Pttime samples (7,781). In the Falkor and MESOSCOPE datasets that were fully labeled by an MS expert, approximately 70% (69% and 73%, respectively) of the features were given a “Bad” designation, corresponding to noise MFs that the expert would not have included in a downstream analysis. In both, 5% of the MFs were unable to be assigned confidently to either “Good” or “Bad” classes and 10% were identified as appearing only in the standards, leaving only ∼15% of the features classified as “Good” (16% and 12%, respectively).

Most metrics had reasonably normal distributions after the scaling and normalization described in Methods. Visually, the most compelling separations between good and bad MFs were observed in our peak shape and novel SNR metrics, with almost complete separation between good and bad peaks provided by the new peak shape metric alone. Peak width and its standard deviation also showed reasonable separation between good and bad MFs (good MFs tended to have low SDs and larger peak widths). The isotope shape and area correlations also showed good separation (Supp. figure 1).

### Logistic regression performance

According to all three logistic regression models (see Methods), the majority of MFs were estimated to have a less than 1% chance of being good. The full model (containing all evaluated peak metrics) and the XCMS model (built on only those metrics calculated from the XCMS output) both displayed a strongly bimodal distribution, with a large number of MFs also exceeding a 99% chance of being good, while the two-parameter model (consisting of the novel SNR metric and the peak shape correlation metric) had a flatter distribution with fewer high-confidence MF assignments and more intermediate values (Figure 1).

**Figure 1:**
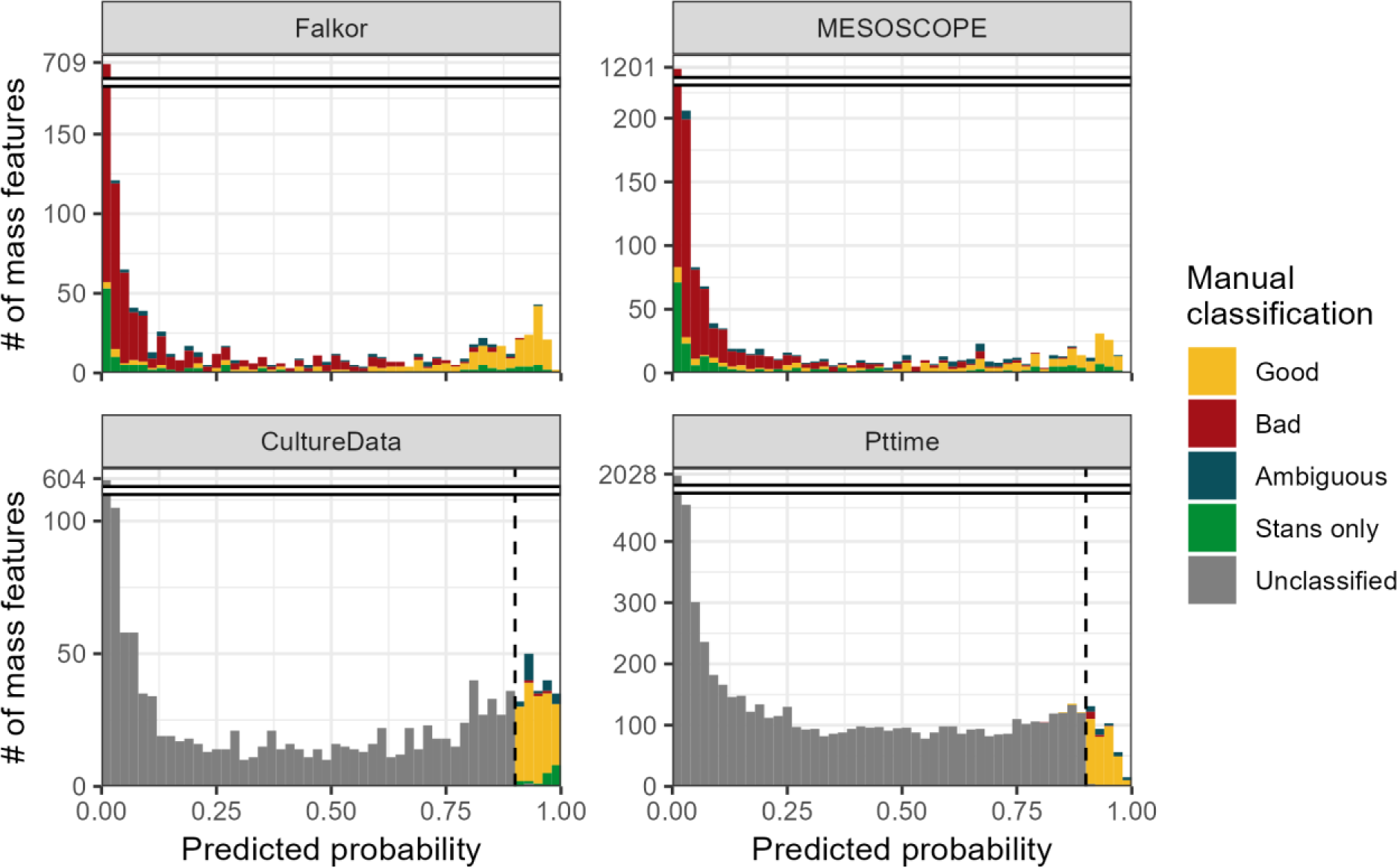
Histograms showing the estimated likelihood of a given mass feature being categorized as “Good” according to the two-parameter logistic model trained on the combined fully-labeled Falkor and MESOSCOPE environmental datasets. Colors indicate the category in which each feature was manually assigned by an expert, with “Stans only” referring to a good mass feature that was only visible in the standards run alongside the samples. Culture datasets CultureData and Pttime were manually labeled only for those features with an estimated likelihood above 90% (dotted black vertical line) according to the final model and were otherwise unclassified.

We explored the relative predictive power of the individual parameters using the full model and found that the predictors least likely to be different from zero due to chance were the mean *m/z* ratio, our novel peak shape correlation metric, and our novel SNR estimate, all with reported p-values < 10^−10^. The value of the novel parameters was then validated using a random forest model that also found them to have the highest importance (Supp.table 1).

The full model performed very well when tested internally on the same dataset both during 80/20 cross validation and when using the full dataset, with FDR (false discovery rate, defined as the number of false positives divided by the total number of positive predictions) values in the 5-10% range and 80-90% GFF (% good features found, defined as the number of true positives divided by the total number of features manually classifed as good) values implying that a large majority of the good MFs passed the threshold with very little noise included. The XCMS metrics performed slightly worse, with FDR values in the 10-15% range and GFF values closer to 75%. The two-parameter model performed worst when tested internally, with an FDR of about 20% and GFF also around 75% (Figure 2). However, when the models were trained on a different dataset than the one they were used to predict classifications for, they all had similar performance with FDRs around 10-25 and GFF around 60-80. The model trained on MESOSCOPE and tested on Falkor had consistently higher values, indicating that it was favoring more MFs recovered at the cost of a higher FDR, while the reverse was true for the model trained on Falkor and tested on MESOSCOPE (Figure 2).

**Figure 2:**
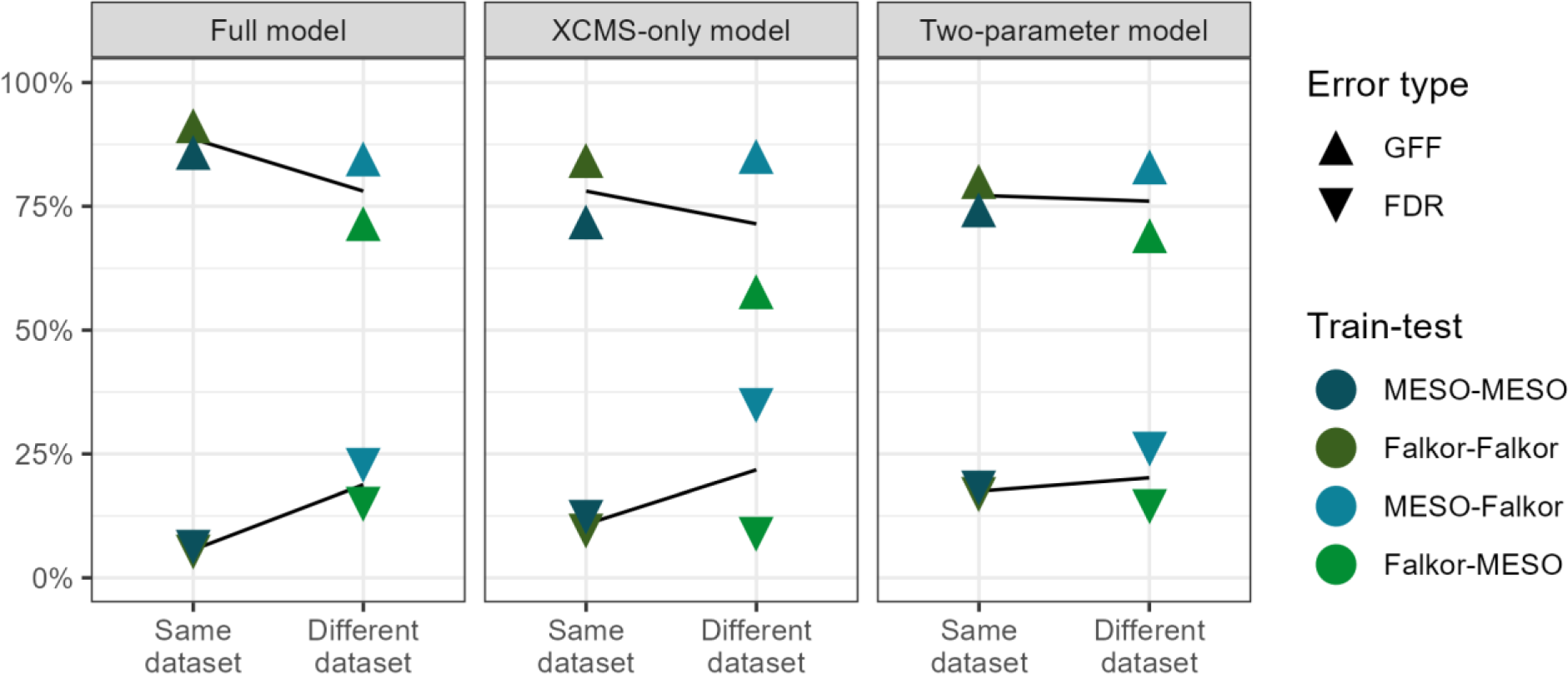
False discovery rate (FDR) and fraction of good features found (GFF) plotted across different subsets of model parameters. Lower FDR indicates a smaller fraction of false positives among those mass features the model categorized as “Good” using a threshold of 0.5, and higher GFF indicates a larger fraction of the total good features were found using the same threshold. Points are colored by the model used for training and testing, with internal validation (using the same dataset for training as prediction) in the darker colors on the left and external validation (using a different dataset for training than prediction) in the lighter colors on the right of each panel. Lines of best fit have been estimated and plotted in black behind the data points, with the steeper slopes found in the full and XCMS-only models indicating overfitting on the training data.

### Model stability under different training sets

We found that the predictions made from a Falkor-trained dataset consistently differed from a MESOSCOPE-trained dataset for the full and XCMS-only models. In the raw probability space, the two-parameter models had the highest Pearson correlation coefficient (*r*) value of 0.996, while the full models and the XCMS-trained models had *r* values of 0.799 and 0.863, respectively. When compared in ranked space using Spearman’s ranked correlation, we found an intensification of this effect, with a higher *ρ* for the two-parameter model of 0.998 but lower *ρ* values for the full and XCMS-trained model of 0.725 and 0.804, respectively (Figure 3).

**Figure 3:**
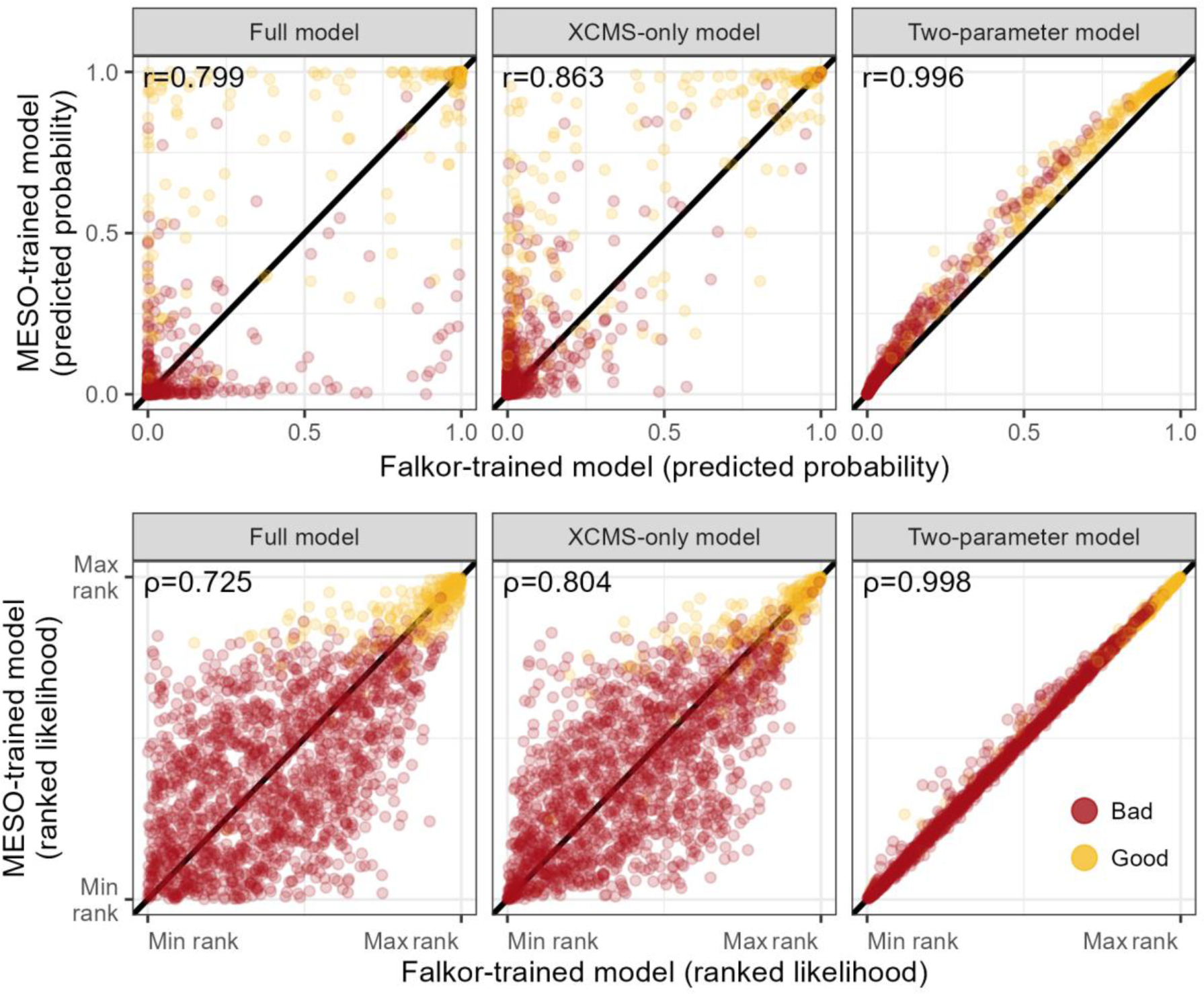
Predicted likelihood of a feature being “Good” according to a model trained on the MESOSCOPE dataset vs a model trained on the Falkor dataset. The top row of plots show the exact likelihood predicted by the logistic model across three different subsets of parameters, while the bottom row shows the estimates ranked from least likely to most likely. Points are colored by their manually-assigned quality according to an expert.

A majority of the time, the estimates from the two models disagreed by more than two times the standard error of the estimate. Some parameters disagreed not only in magnitude but also in sign, with the Falkor-trained full model increasing MF goodness likelihood with larger PPM variation and a wider peak width, while the MESOSCOPE-trained full model had negative estimates for each of these parameters. Notably, the peak shape and novel SNR parameters used in the two-parameter model were among the most robust to training model variation, potentially explaining the consistency described above (Supp. figure 2).

When testing model stability under a smaller sample size, we found reasonably good convergence in a dataset containing half the mass features with most model parameters falling within two standard errors of the estimate for the XCMS and two-parameter model, while the full model required closer to 80% of the mass features to produce estimates consistent with the original model (Supp. figure 3).

### Regularized regression and random forests perform about the same

None of the penalized regression models significantly improved cross-validated performance between the MESOSCOPE and Falkor datasets when measured by both initial performance and the performance drop when applied across datasets. All three regularized regression models had similar behavior, with ridge regression (α = 0) obtaining the lowest rates for both GFF and FDR, while lasso (α = 1) obtained higher rates for both and represented a less-stringent false negative acceptance. As expected, the elastic net (α = 0.5) fell in between the two (Supp. figure 4). The random forest model, interestingly, had perfect predictive capacity when tested internally on the training data (FDR=0%, GFF=100%, for both MESOSCOPE and Falkor) but showed a significant drop in improvement when applied across datasets (Supp. figure 4). In each case, the performance drop when applied to a novel dataset was more extreme than the simple two-parameter model described above.

### Performance of a stricter threshold on novel datasets

We settled on a 90% likelihood threshold for application to novel datasets because it struck a balance between the number of MFs we estimated to be necessary for robust testing while still remaining reasonable to manually label. For the CultureData dataset, we obtained 1,790 total mass features, 192 of which had predicted likelihoods above 0.9. Of these, 151 were identified manually as “Good”, 21 were given “Ambiguous” designations, and only 3 were flagged as “Bad”, with the remaining 17 appearing only in the standards. For the Pttime dataset, 7,781 were obtained with 400 flagged by the model as “Good”. 348 were truly good MFs, 35 were ambiguous, and 17 were “Bad”. No standards were run during this analysis, so there were no features in that category.

With the stricter threshold, we obtained FDR values consistently below 5% even on the novel datasets, with values of 1.0%, 0.0% (truly zero false positives), 2.0%, and 4.6% for Falkor, MESOSCOPE, CultureData, and Pttime respectively (Figure 4). Of course, this low error rate meant that we miss out on additional potentially valuable features, with only a fraction of the total good MFs making it past this threshold. In both the Falkor and MESOSCOPE datasets, less than half of the good MFs were labeled as such, with actual values of 39.4% and 26.5%, respectively. Since we did not label the complete dataset for CultureData and Pttime, we cannot accurately calculate the GFF but expect it to be in a similar range (Figure 4).

**Figure 4:**
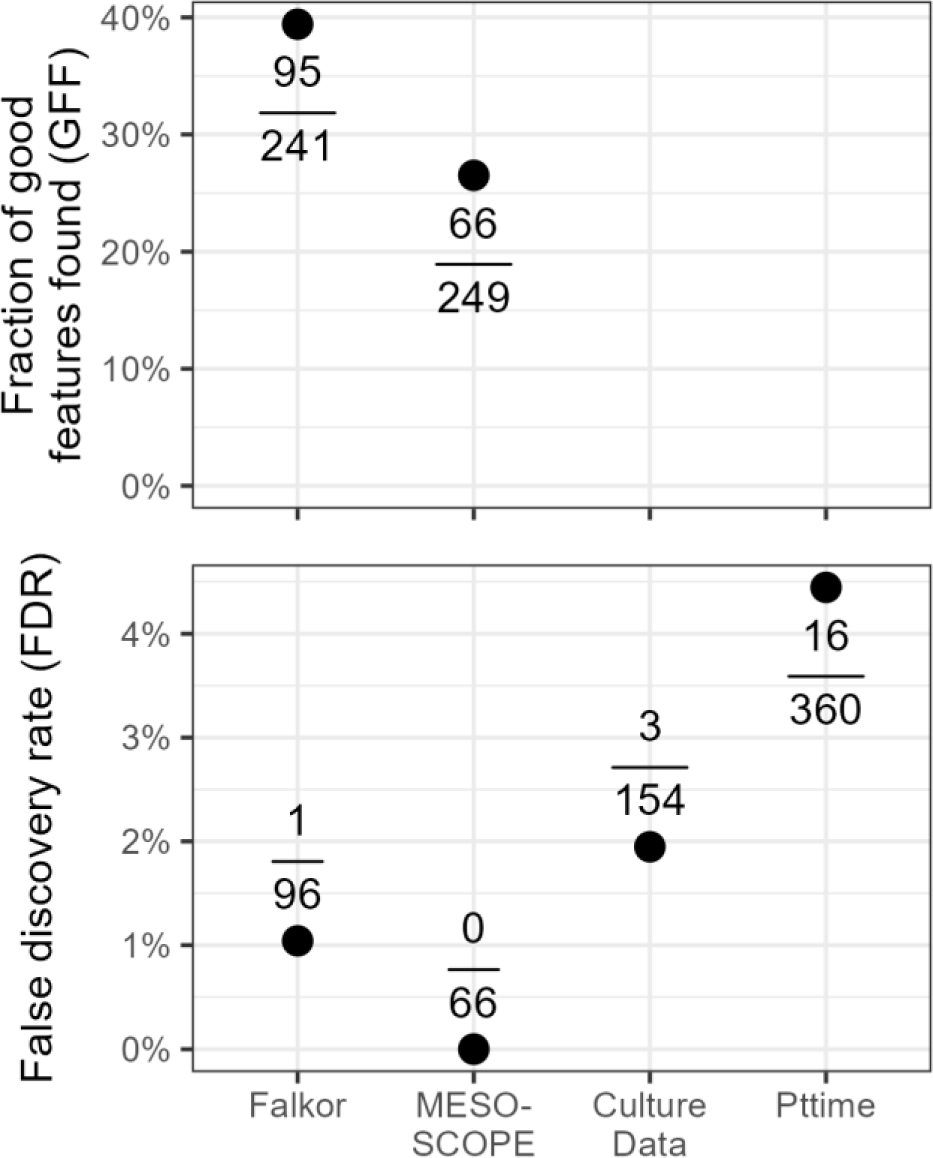
False discovery rate and proportion of total good features identified as good by the two-parameter model trained on the combined MESOSCOPE/Falkor dataset and applied to each dataset. A stricter likelihood threshold is used here (0.9) than in Figure 2. FDR is calculated by dividing the number of false positives by the total positives produced by the model and GFF is calculated by dividing the number of true positives by the total number of good features as identified manually (only possible for fully-labeled datasets). Points correspond to the calculated percentage and absolute numbers are provided above/below the point.

### Implications for biological conclusions

#### Univariate techniques

A majority of the features (1,323 of 1,832 total) in the original, non-thresholded MESOSCOPE dataset had no significant trend with depth, with FDR-controlled Kruskal-Wallis p-values exceeding 0.05 (Figure 5). The largest category that did have a trend with depth was the 15m = DCM > 175m category, containing 118 mass features, with largest peak areas distributed evenly between the 15 meter and deep chlorophyll maximum (DCM) samples, while the 175 meter samples had significantly smaller areas. The similar but statistically distinct categories of 15m > DCM > 175m and DCM > 15m > 175m had 68 and 35 features, respectively, and together indicate that many molecules are highly abundant throughout the surface ocean down to the DCM layer and decrease in concentration at 175 meters. A surprising number of features were also found to have DCM minima (DCM < 15m = 175m, 26 features) or linear increases with depth (15m < DCM < 175m, 12 features) given the few environmental parameters that have these trends (Figure 7).

**Figure 5:**
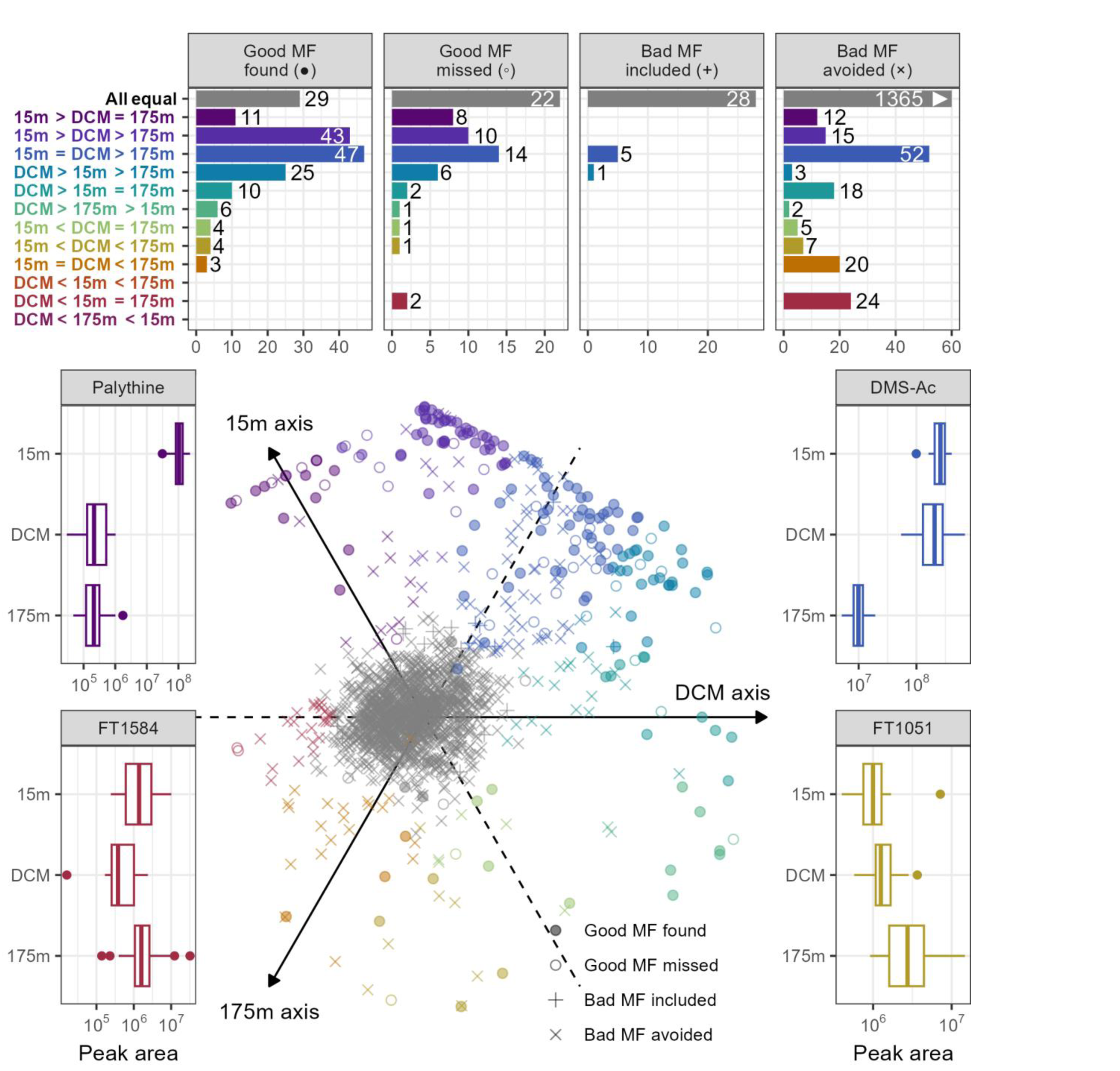
Plot of metabolite response to depth and the model classification error distribution. Barplots at the top show the number of MFs (mass features) in each depth response category and are broken down by the classification error types. Compounds were assigned a depth category via Dunn’s post-hoc test for significant differences between the sample depths. Good MF found = true positive, good MF missed = false negative, bad MF found = false positive, and bad MF avoided = true negative according to a 0.5 likelihood threshold. Note that the majority of the features in the bad MF avoided category fell into the “All equal” depth class for which there was no significant differences between the depths (1365 MFs) but the x-axis has been truncated at 60 for ease of visualization. The boxplots in the bottom illustrate the depth response type for 4 specific categories, with raw peak area plotted on a log scale against the sample depth (DCM = deep chlorophyll maximum, ∼110 meters). All MFs are shown in the central bottom plot across three axes using the rank-normalized median value at each depth as the coordinate for that axis. Each mass feature corresponds to a point in the plot, and their position on the plot describes the shape of their depth profile. Compounds aligning with the 15m axis correspond to compounds with most of their abundance found in the surface ocean; points far to the right side correspond to compounds that are found only at the DCM; points found at the bottom of the plot are those compounds that increased more or less linearly with depth.

A different story emerged, however, when the bad MFs were removed from this analysis. Good features were most commonly found to have their highest concentrations at the DCM or the surface, rather than being fixed with respect to depth. Of the 182 good MFs, less than a fifth had no trend with depth (44/249) and a majority had unequivocally lowest values in the 175 meter samples (those with 15m/DCM > 175m, 145 features). The two-parameter model, when applied with a 50% likelihood threshold, also recovered this general feature distribution and classified many of the features with no significant depth signal as likely to be bad.

Additionally, a large number of features manually identified as bad nonetheless had significant differences with depth. This was surprising because we had assumed that bad MFs corresponded to instrument or chemical noise, which we did not expect to have any biological trend. Further investigation of a few randomly selected bad features with a biological difference revealed the reason behind this: most of those investigated were actually tails of other MFs. Integrating just the tail of a peak retains the biological signal of the full peak while still looking visually like instrument noise, thereby introducing pseudoreplication in the feature space.

The model did fail to recover some interesting biological variation, however. Two features of particular interest were those good MFs with a DCM minimum (DCM < 15m = 175m), both of which were missed by the two-parameter model. These features possess an unexpected biological signal that does not track with depth or other common oceanographic parameters, thereby potentially representing an interesting biomarker that decreases despite an increase in biomass.

#### Multivariate techniques

Multivariate statistics also benefitted from the reduced FDR when applying the two-parameter model. For the PERMANOVAs, we found that the proportion of variance explained (R^2^) and the pseudo-F statistic increased monotonically with the likelihood threshold used to subset the data (Table 1). In each test, the permutational p-value obtained was less than 0.001, indicating that the differences between samples due to depth were unlikely to be due to chance. However, the pseudo-F and was much larger with higher thresholds, scaling from around 8.5 when thresholding at a 1% likelihood to 42 when thresholded at a 90% likelihood (Table 1).

**Table 1:** Number of mass features, percent variance explained, pseudo-F statistic, and stress values from performing a permutational MANOVA and 2D non-metric multidimensional scaling (NMDS) on subsets of the full mass feature selection according to variable likelihood thresholds.

| Data subset | # of MFs | % Var Expl. | Pseudo-F | 2D NMDS stress |
| --- | --- | --- | --- | --- |
| All MFs | 2086 | 10.4 | 5.5 | 0.169 |
| Threshold 0.01 | 1129 | 15.2 | 8.6 | 0.191 |
| Threshold 0.1 | 516 | 25.0 | 16.0 | 0.200 |
| Threshold 0.5 | 287 | 34.6 | 25.3 | 0.180 |
| Threshold 0.9 | 75 | 46.7 | 42.0 | 0.097 |
| Only good MFs | 249 | 44.2 | 38.1 | 0.142 |

We also tested the inclusion of all the features identified with XCMS (corresponding to a 0% threshold) and the results when only the manually-identified “Good” features were included. The default XCMS output continued the trend observed above, as expected, with the least variance explained and the lowest R^2^ value. Subsetting for the “Good” MFs only, however, did not actually return the highest F-ratio or R^2^, instead falling between the 50% and 90% thresholds for these two metrics. In large part this is due to the much smaller number of features: 249 features were manually labeled as Good, while only 75 exceeded the 90% likelihood threshold (Table 1).

The relative power of identifying only the very best MFs was also illustrated visually with non-metric multidimensional scaling (NMDS) plots (Figure 6). In these common exploratory plots, the MFs with likelihoods above 50% strongly separated by depth while lower thresholds disguised the true signal and had higher stress values. Performing an NMDS on the manually-identified Good MFs resulted in output nearly indistinguishable from those of the 90% and 50% thresholds.

**Figure 6:**
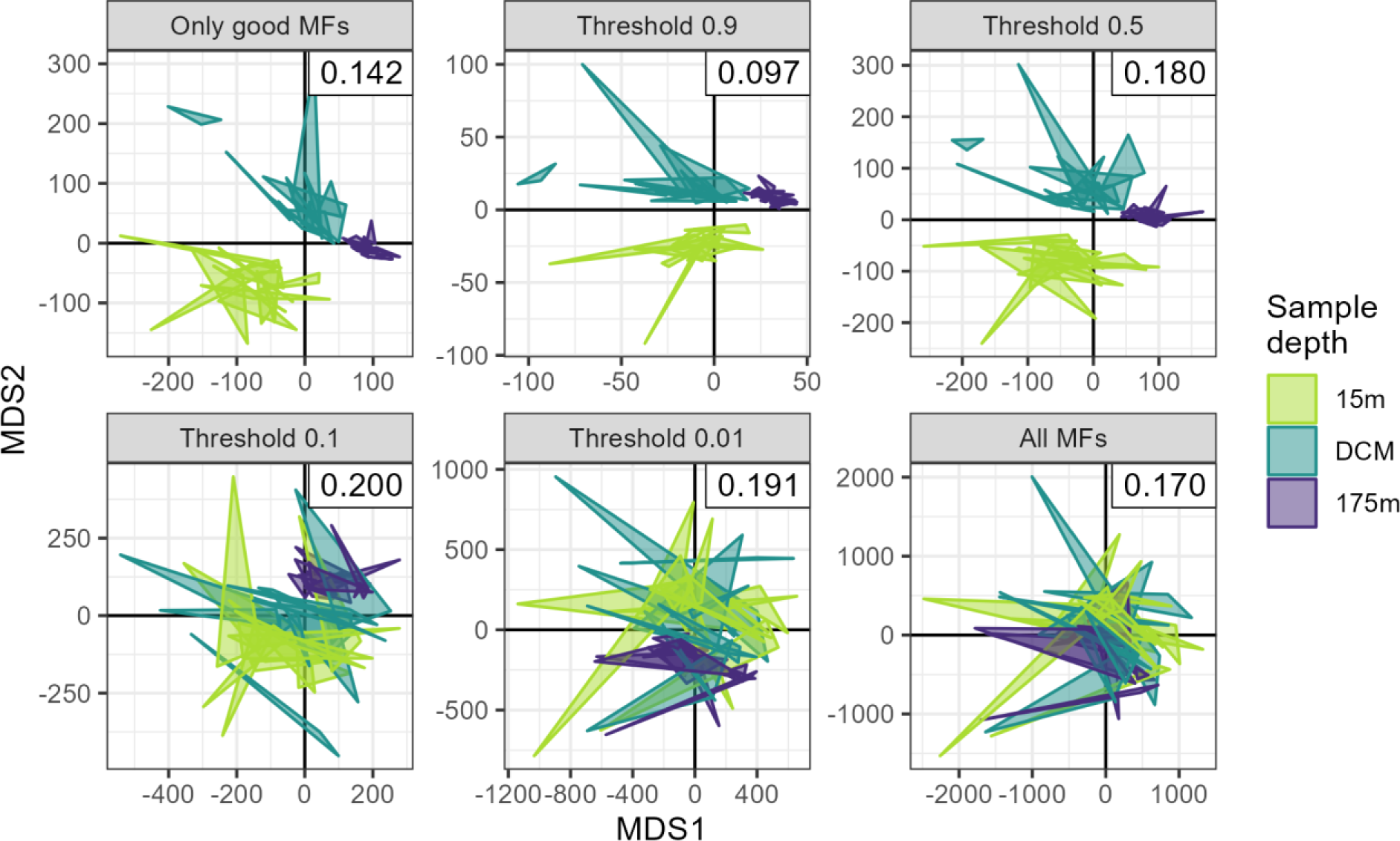
2D non-metric multidimensional scaling (NMDS) plots of metabolite similarity according to sample depth across multiple likelihood thresholds. Triplicate samples are represented by the vertices of the triangles and colored by the depth from which they were sampled (DCM = deep chlorophyll maximum, ∼110m). “Only good MFs” refers to those features manually labeled as “Good”. NMDS stress values are reported in the upper right corner of each plot.

## Discussion

We used two fully-labeled and two partially-labeled HILIC LC-MS datasets to assess the performance of the XCMS algorithm and construct a robust model of peak quality. To measure performance, we used two measurements of success closely tied to intuitive questions about a dataset: the percentage of total good features found (GFF, also known as recall or sensitivity) and the percentage of bad mass features (MFs) included, also known as the false discovery rate (FDR). We decided against using the F_1_ score as an overall summary statistic because false negatives and false positives have very different implications in this context and should be treated separately.

One of the major ways in which this manuscript differs from prior work is its focus on summary statistics calculated across multiple files. Most existing peakpicking literature uses the single-file EIC peak as the core bit of training data, but that approach ignores critical information obtained elsewhere in the MS run that can change the judgement made on a single chromatogram (Pirttilä et al. 2022; Guo et al. 2021; Müller et al. 2020). Features that are high quality are typically represented in multiple files, especially in quality control pooled samples, and a feature that only has a peak in a single file is typically regarded as highly suspect, if not removed entirely. Classifying an entire feature at once not only has the advantage of reducing the amount of manual labor by a factor equal to the number of files in the run (typically 10s-100s smaller) but is also a better representation of the judgement made by an MS expert. An exemplary implementation of this multi-file approach in prior work is reported in Kantz et al. (2019), who compared the multi-file summary statistic model to a deep neural network and came to many similar conclusions.

### Two-parameter logistic regression model with raw data metrics showed the most reliable performance

We explored several different types of classification models for separating good mass features (MFs) from bad, with a particular focus on quantifying the likelihood of each class rather than just returning the label. We found that a simple two-parameter logistic regression model trained on two novel metrics of peak quality had reasonably good performance on the training set and was highly robust when applied to novel datasets. The logistic regression in particular was favored over the random forest and regularized regression we tried due to their similar performance and increased interpretability (Supp. figure 4).

This model outperforms the previously reported logistic regression model in Kantz et al. (2019) and is highly simplified. There, they used a nineteen-parameter multiple logistic regression model and found a maximum performance of 80% GFF and an FDR of 34% on a cross-validated second cohort, similar to our cross-dataset testing. Our final two-parameter model also had an 80% GFF at a 0.5 likelihood threshold, but a significantly lower FDR of ∼22.9%. This increased performance is likely due to our use of the metrics recalculated from the raw data, as their metrics were only calculated from the default MzMine2 peak parameters reported: similar to our XCMS-only model. Previous work on a “shape-orientated” algorithm also established the utility of testing the extracted ion chromatogram against a Gaussian shape (Bai et al. 2022). There, the use of a Marr wavelet had GFF values in the 98-100% range but very high FDR values of 82-91%, representing a very lenient threshold much closer to the XCMS or ADAP defaults.

#### Performance relative to recent deep learning methods

Guo et al. (2021) presented EVA and reported an accuracy of 90-95%, a range inclusive of our accuracy on both the Falkor (92.1%) and MESOSCOPE (94.4%) datasets when using a likelihood threshold of 0.5. However, we note that accuracy alone can be a highly misleading statistic to report when working with unbalanced datasets because very high accuracy can be obtained by simply classifying everything as bad, with a strong incentive to actually *increase* the number of bad MFs initially picked while doing so. This strategy, when applied to our data, returned accuracies in the 80-90% range despite being a useless classifier for downstream analysis.

The class imbalance, with mostly poor quality MFs, is partially why we chose to measure precision and recall instead of total accuracy. However, precision and recall can also be ambiguous when the positive class is not specified and the raw confusion matrices are unavailable, thus our very precise use of the FDR and GFF metrics as well as providing the confusion matrices in Supplemental Table 2. Melnikov, Tsentalovich, and Yanshole (2020) reported precision and recall in their presentation of peakonly, relative to which we obtained higher accuracy (they report 89% accuracy) but worse GFF and FDR (89% and 3%, respectively, relative to our 77.1% GFF and 19.6% FDR overall). However, if we report precision and recall with the positive class set to “Bad”, essentially trying to predict poor-quality MFs instead of good ones, our precision becomes 96.5% and our recall 95.7% due to the strong prior information about most MFs being bad.

Gloaguen, Kirwan, and Beule (2022) later introduced NeatMS, another CNN, and compared it directly to peakonly to claim equivalent or superior performance across a range of dilution factors. However, they do not report total precision or recall metrics in a comprehensive untargeted way but instead focus only on assessing the model’s performance on known chemical standards. They report a percentage of standards found for the peakonly model applied to their data and find that its performance is significantly lower (79.4%) than the recall reported in Melnikov, Tsentalovich, and Yanshole (2020), perhaps indicating that the peakonly model is still overfit.

While the model we present here likely has reduced performance relative to the CNNs, we would argue that its utility is not in maximizing performance but instead in maximizing interpretability, as previously noted by Kantz et al. (2019). In particular, the CNNs provide no way to control the tradeoff between false positives and false negatives and no relative ranking of individual MF quality beyond the broad bins into which they are placed or explanation of relative metric strength for later analyses.

#### Assessing the relative power of individual metrics

Although the deep learning models show promise for peak quality recalibration, many mass-spectrometrists are reluctant to jump fully to their black-box nature. For this reason, we also reported here the relative power of individual parameters in our full model and use the results to dispel several myths about which parameters are useful in distinguishing signal from noise.

The two metrics in the final model were rederived from the raw EIC data because they matched our intuition about what makes an MF look good to an MS expert. These are very simple metrics and therefore fast to calculate, but we expect that more complicated metrics could perform even better. For example, the method of using the data within the peak boundaries for SNR calculation rather than data outside of them is not known to the authors to be implemented elsewhere but could be further improved by more advanced smoothing methods rather than using the residuals directly. Additionally, the calculation of peak shape using a Pearson’s correlation to an idealized curve was not expected to be especially powerful given prior research (e.g. Ipsen et al. (2010)) and that the centWave algorithm essentially uses this information already during the wavelet fitting, but still proved to be a highly informative parameter. This metric could be improved with more careful summary statistics that account for the differences between samples. Currently, the use of the overall median value does a reasonable job at identifying MFs that appear in many samples but performs poorly when detecting MFs that appear in only a few. Also worth noting is that the calculation of any new metrics such as these that rely on access to the raw data require exact specification of the maximum and minimum *m/z* and retention time for a peak, values that are not always returned by peakpicking algorithms and must be recalculated, as in Kantz et al. (2019). To avoid the additional overhead of recalculation and the possiblity of raw data unavailability, we have implemented these metrics during the initial peakpicking step of XCMS in a fork of the GitHub available at https://github.com/wkumler/xcms and have submitted a pull request to implement them into XCMS directly.

We were surprised at the poor performance of several other metrics. The isotope information in particular was expected to be a very strong predictor of MF quality given previous work that uses this metric extensively (Libiseller et al. 2015; Treutler and Neumann 2016; El Abiead et al. 2021). We learned that many noise MFs still have reliable isotopes (perhaps unsurprising, given that the noise is in fact often caused by solvents or contaminants that are still chemical in nature) and that many real MFs are simply too small (low-intensity) to have detectable isotope peaks in this kind of dilute environmental sample.

The relative standard deviation (RSD), also called the coefficient of variance, among pooled samples is another parameter that performed surprisingly poorly given its general acceptance as a quality scoring metric. In the full model, neither the traditional calculation of RSD (standard deviation divided by the mean) nor the robust implementation (median absolute deviation divided by the median) were significant parameters. This result was also reported by Gloaguen, Kirwan, and Beule (2022) who noted that while the RSD was typically lower for high-quality features there were many noise MFs with low RSDs as well.

We also showed that the automatically calculated SNR parameter from XCMS is not especially useful in distinguishing signal from noise. After inspecting a selection of MFs that had anomalous values for this metric, we are inclined to agree with Myers et al. (2017a) and conclude that this is often due to insufficient data outside of the peak for a robust calculation of noise level.

Finally, we were surprised to find essentially no predictive power offered by peak area or intensity, with good MFs distributed almost identically to the bad MFs in this space. This cautions strongly against an arbitrarily-decided intensity threshold for winnowing down the number of MFs, in agreement with previous work (Houriet et al. 2022; Barupal et al. 2021). Similarly surprising was the lack of power in the design-of-experiments metrics, although this was less surprising given the number of missing values that were later filled in with an order of magnitude outside the most extreme value (Supp. figure 1).

#### Model selection and simplification

We settled on the highly reduced model of just two parameters because we found that additional parameters often improved performance on the training set but did not do so significantly for the novel datasets where the application of such a model is actually useful (Figure 2). The drastic drop in performance on out-of-sample data was particularly concerning because it creates overconfidence in the true level of noise actually ending up in the final dataset. One important caveat to note is that for the partially-labelled CultureData and Pttime datasets, there exists an uncontrolled degree of experimenter bias because the MS expert responsible for labeling did know that these MFs were all expected to be good. However, given that we do still see poor-quality MFs in this set indicates that this was not an overwhelming bias.

We also found that this reduced two-parameter model was largely independent of the particular training set used, unlike in the more complex models (Figure 3). This was true in both absolute likelihood as well as rank-ordered space, a particularly important distinction when one imagines manually labeling “down” the dataset where the researcher starts viewing the chromatograms associated with the very best features and eventually reaches a point where enough MFs have been reviewed or bad MFs are frequent enough that they decide to stop.

A final benefit to the reduced model is the smaller training set required to reach stability (Supp. figure 3). This reduced size means that a useful model could be trained using only a fraction of the MFs identified in a sample set and then used to predict the quality for the remainder of the features. This reduction in training set size was not as significant as we expected, however, with several hundred features requiring manual labeling before even the two most stable parameters reached a consensus.

### Biological conclusions vary significantly by feature quality

We found that the conclusions obtained from the metabolomic datasets differed in significant ways depending on the quality threshold used to remove bad MFs from the downstream analysis. In the multivariate case, we ran the same analysis of PERMANOVAs and NMDS plots on various subsets of the original XCMS output and found that the effect size of depth was strongly influenced by by the threshold chosen. This is unsurprising given that most noise MFs should not have a biological signal to begin with, but is troubling for interpreting analyses where the FDR is not reported or the dataset not manually reviewed because the absence of a notable effect could simply be due to the overwhelming degree of noise in the default output.

In the univariate case, we showed that while noise MFs are predominantly absent of a large biological signal, there are many that still have a significant biological trend. While some of these are inherently due to the likelihood of getting a small p-value with enough attempts despite FDR correction, a larger number of these poor-quality MFs were due to partial integration in which only the tail of a feature was integrated. This essentially duplicates the signal of the original MF in later analyses and should be removed. The real features showed a strong biological trend of high concentration throughout the surface ocean and down through the deep chlorophyll maximum (DCM), with most features equally abundant at 15 meters and this ∼110 meter depth feature before dropping off at depth. This pattern tracks well with previous reports of biomass from the same sample site as well as earlier literature (Barone et al. 2022; Heal et al. 2021). Critically, this also highlights the danger of noise MFs when additional normalizations are later applied. Scaling metabolomic data to biomass measurements is a common technique, and yet here it would have caused an enormous number of false positives that would have appeared to be intriguingly enriched below the DCM.

### Conclusions

The large number of mass features due to noise present in metabolomics datasets can be controlled using a simple logistic classification model. We trained such a model on two full-labeled open ocean HILIC datasets and found that the best performing parameters in the model were a custom signal-to-noise metric and a test of similarity to a bell curve. This model showed robustness to overfitting, independence from the training set, and a reduced degree of manual labeling required. With this model, we showed how the distribution of metabolites in the open ocean is strongly affected by depth and categorized molecules according to their depth response. This distribution reproduces measures of bulk biomass but highlights several molecules of interest that diverge from the overall trend.

## Methods

### Sample collection

Environmental samples were collected from the North Pacific Subtropical Gyre near Station ALOHA during two research cruises that targeted strong mesoscale eddy features during June/July 2017 and March/April 2018, traversing an area between 28 °N, 156 °W and 23 °N, 161 °W. An eddy dipole off the coast of Hawaii was detected using sea-level anomaly (SLA) satellite data and targeted for both a transect across the cyclonic and anticyclonic poles of the eddy dipole. The cyclonic pole of the eddy had a maximum negative SLA anomaly of −15 cm in 2017 and −20 cm in 2018, while the anticyclonic center reached +24 cm in 2017 and +21 cm in 2018. The 2017 cruise samples were taken along a transect across the eddy dipole while the 208 cruise targeted only the center of each eddy.

Environmental samples were obtained using the onboard CTD rosette to collect water from 15 meters, the deep chlorophyll maximum (DCM), and 175 meters during the 2017 MESOSCOPE cruise and from 25 meters and the DCM during the 2018 Falkor cruise. The DCM was determined visually from fluorometer data during the CTD downcast and Niskin bottles were tripped during the return trip to the surface. Seawater from each depth was sampled in triplicate by firing one Niskin bottle for each sample. Samples were brought to the surface and decanted into prewashed (3x with DI, 3x with sampled seawater) polycarbonate bottles for filtration. Samples were filtered by peristaltic pump onto 142mm 0.2 µm Durapore filters held by polycarbonate filter holders on a Masterflex tubing line.

Pressures were kept as low as possible while still producing a reasonable rate of flow through the filter, approximately 250-500 mL per minute. Samples were then removed from the filter holder using solvent-washed tweezers and placed into pre-combusted aluminum foil packets that were then flash-frozen in liquid nitrogen before being stored at −80 °C until extraction. A methodological blank was also collected by running filtrate through a new filter and then treated identically to the samples.

Culture samples used as the validation sets for this paper have been previously described by Durham et al. (2022) and on Metabolomics Workbench (Project ID PR001317).

### Sample processing

Extraction of the environmental samples followed a modified Bligh & Dyer approach as detailed in Boysen et al. (2018). Briefly, filters were added to PTFE centrifuge tubes with a 1:1 mix of 100 µm and 400 µm silica beads, approximately 2mL −20 °C Optima-grade DCM, and approximately 3mL −20 °C 1:1 methanol/water solution (both also Optima-grade).

Extraction standards were added during this step. The samples were then bead-beaten three times, followed by triplicate washes with fresh methanol/water mixture. Samples were then dried down under clean nitrogen gas and warmed using a Fisher-Scientific Reacti-Therm module. Dried aqueous fractions were re-dissolved in 380 µL of Optima-grade water and amended with 20 µL isotope-labeled injection standards. Additional internal standards were added at this point to measure the variability introduced by chromatography and ionization, and the reconstituted fraction was syringe-filtered to remove any potential clogging material. This aqueous fraction was then aliquoted into an HPLC vial for injection on the HILIC column and diluted 1:1 with Optima-grade water. A pooled sample was created by combining 20 µL of each sample into the same HPLC vial, and a 1:1 dilution with water half-strength sample was aliquot from that to assess matrix effects and obscuring variation (Boysen et al. 2018). Also run alongside the environmental samples were two mixes of authentic standards in water and in an aliquot of the pooled sample at a variety of concentrations for quality control, annotation, and absolute concentration calculations. HPLC vials containing the samples were frozen at −80 °C until thawing shortly before injection.

The CultureData samples were re-run from the frozen aliquots for this paper. The Pttime sample processing is documented on Metabolomics Workbench where it has been assigned Project ID PR001317.

### LC conditions

For the MESOSCOPE, Falkor, and CultureData samples a SeQuant ZIC-pHILIC column (5 um particle size, 2.1 mm x 150 mm, from Millipore) was used with 10 mM ammonium carbonate in 85:15 acetonitrile to water (Solvent A) and 10 mM ammonium carbonate in 85:15 water to acetonitrile (Solvent B) at a flow rate of 0.15 mL/min. The column was held at 100% A for 2 minutes, ramped to 64% B over 18 minutes, ramped to 100% B over 1 minute, held at 100% B for 5 minutes, and equilibrated at 100% A for 25 minutes (50 minutes total). The column was maintained at 30 °C. The injection volume was 2 µL for samples and standard mixes. When starting a batch, the column was equilibrated at the starting conditions for at least 30 minutes. To improve the performance of the HILIC column, we maintained the same injection volume, kept the instrument running water blanks between samples as necessary, and injected standards in a representative matrix (the pooled sample) in addition to standards in water. After each batch, the column was flushed with 10 mM ammonium carbonate in 85:15 water to acetonitrile for 20 to 30 minutes. LC conditions for the Pttime samples are documented on Metabolomics Workbench where it has been assigned Project ID PR001317.

### MS conditions

Environmental metabolomic data was collected on a Thermo Q Exactive HF hybrid Orbitrap (QE) mass spectrometer. The capillary and auxiliary gas heater temperatures were maintained at 320°C and 100°C, respectively. The S-lens RF level was kept at 65, the H-ESI voltage was set to 3.3 kV and sheath gas, auxiliary gas, and sweep gas flow rates were set at 16, 3, and 1, respectively. Polarity switching was used with a scan range of 60 to 900 m/z and a resolution of 60,000. Calibration was performed every 3-4 days at a target mass of 200 m/z. DDA data was collected from the pooled samples for high-confidence annotation of knowns and unknowns. All files were then converted to an open-source mzML format and centroided via Proteowizard’s msConvert tool. For the Pttime samples, files were pulled directly from Metabolomics Workbench via Project ID PR001317 and used in their existing mzXML format.

### Peakpicking, alignment, and grouping with XCMS

The R package XCMS was used to perform peakpicking, retention time correction, and peak correspondence (Smith et al. 2006; Tautenhahn, Böttcher, and Neumann 2008). Files were loaded and run separately for each dataset using the “OnDiskMSnExp” infrastructure. Default parameters for the CentWave peakpicking algorithm were used except for: ppm, which was set to 5; peakwidth, which was widened to 20-80 seconds; prefilter, for which the intensity threshold was raised to 10^6^; and integrate, which was set to 2 instead of 1. snthresh was set to zero because there are known issues with background estimation in this algorithm (Myers et al. 2017a), and both verboseColumns and the extendLengthMSW parameter were set to TRUE. For retention time correction, the Obiwarp method was used except for the CultureData dataset, which was visually inspected and determined not to require correction (Benton, Want, and Ebbels 2010). For the Obiwarp algorithm, the binsize was reduced to 0.1 but all other parameters were left at their defaults or equivalents.

Peak grouping was performed on the two environmental datasets and the Pttime data with a bandwidth of 12, a minFraction of 0.1, binSize of 0.001, and minSamples of 2 but otherwise default arguments. CultureData’s minFraction was raised to 0.4 but was otherwise identical. Sample groups were constructed to consist of the biological replicates for all datasets. After peak grouping, peak filling was performed using the fillChromPeaks function with the ppm parameter set to 2.5. Finally, mass features with a retention time less than 30 seconds or larger than 20 minutes were removed to avoid interference from the initial and final solvent washes.

### Manual inspection and classification

After the full XCMS workflow was completed, the mass features were visually inspected by a single qualified MS expert. For the Falkor and MESOSCOPE datasets, every mass feature was inspected, while only those features with a predicted probability of 0.9 or higher according to the two-parameter model produced below were inspected for the CultureData and Pttime datasets. Inspection consisted of plotting the raw intensity values against the corrected retention-time values for all data points within the *m/z* by RT bounding box determined by the most extreme values for the given feature. For this step, we decided to plot the entire feature across all files simultaneously rather than viewing each sample individually to both accelerate labeling and to more accurately represent what MS experts typically do when assessing the quality of a given mass feature (Figure 7). We also decided to ignore missing values and linearly interpolate between known data points rather than filling with zeroes. These EICs were then shown to an MS expert for classification into one of 4 categories: Good, Bad, Ambiguous, or Stans only if the feature appeared to only show up in the standards. A few randomly-chosen features from the manually-assigned Good and Bad classifications are shown in Figure 7.

**Figure 7:**
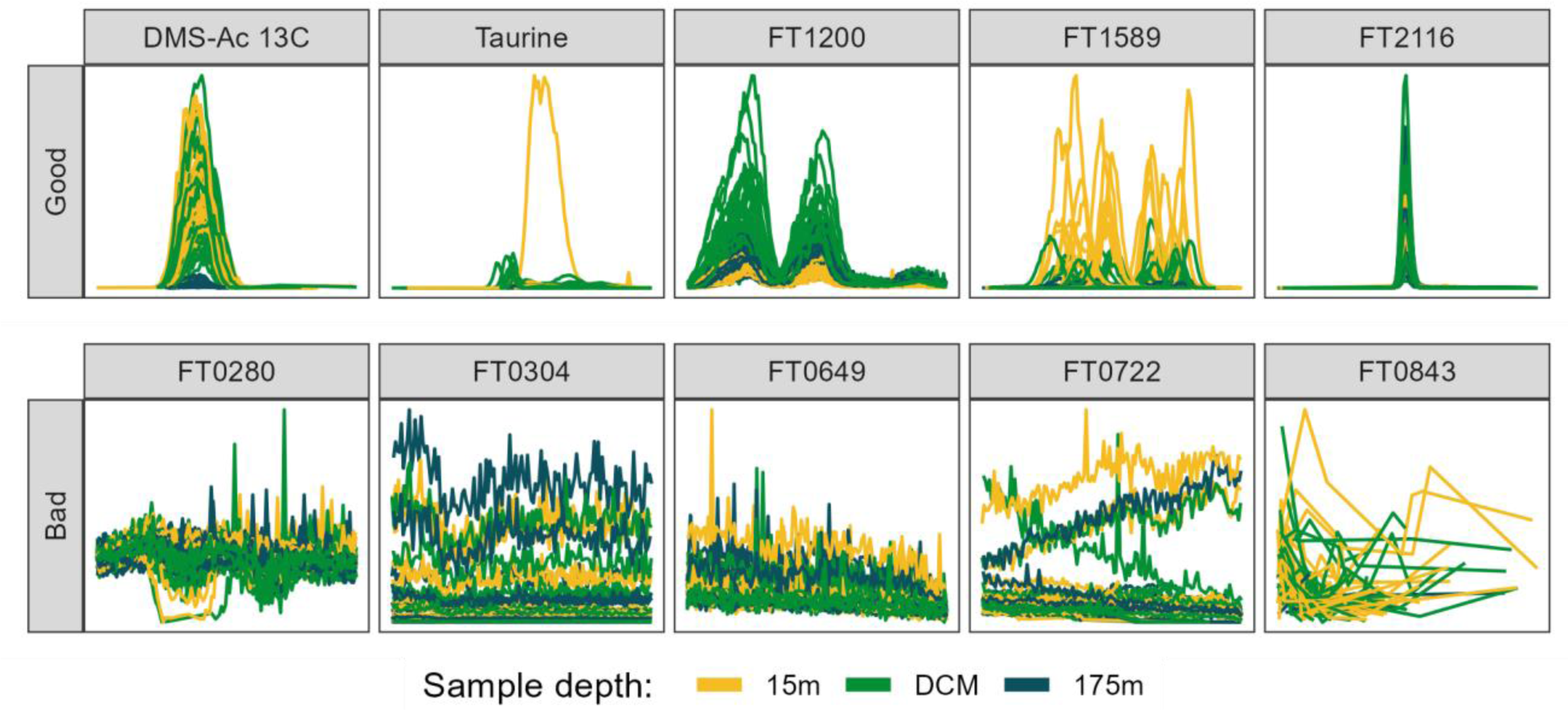
Randomly selected ion chromatograms from both “Good” (top row) and “Bad” (bottom row) manual classifications. Plots show retention time along the x-axis in a 1 minute window around the center of the feature and show measured intensity on the y. Features are from the MESOSCOPE dataset and colored by the depth from which the biological sample was taken. DCM = deep chlorophyll maximum, approximately 110 meters. Mass feature identifications are provided as the title of each panel, starting with “FT” and followed by 4 digits except for the two features annotated using authentic standards run alongside: the 13C isotope of dimethylsulfonioacetate (DMS-Ac) and taurine.

### Peak feature extraction and metric calculation

Our process of feature engineering involved querying several MS experts in our lab about their intuition for what they thought best distinguished poor-quality MFs and noise from good ones. The simplest metrics to calculate were summary statistics of those parameters reported directly by XCMS. These features consisted of the mean retention time (RT) of each MF and the standard deviation (SD) within the feature and the mean peak width (calculated by subtracting the max RT from the minimum) and its SD. We also calculated the mean *m/z* ratio and the SD in parts-per-million (PPM) by dividing each peak’s reported *m/z* ratio by the *m/z* ratio of the feature as a whole, then multiplying by one million. Mean peak area was calculated by taking the log_10_ of the individual areas then taking the mean, and the same process (log_10_ then mean) was repeated for the SD of the peak areas. XCMS’s default signal-to-noise parameter, sn, was also summarized in this way, but we only used sn values that were greater than or equal to zero and replaced any zeros with ones to avoid negative infinities after taking the log_10_. We also used the mean of other parameters reported by XCMS (f, scale, and lmin) as features. We additionally calculated several design-of-experiments metrics, using the number of peaks in each feature divided by the total number of files as well as the fraction of files in which a peak initially found by the peakpicker. This last metric was further subset into the fraction of samples in which a peak was initially found and the fraction of standards in which a peak was found (for those datasets in which standards were run). Finally, the coefficient of variance was estimated for the pooled sample peak areas by dividing the SD of the pooled sample peak areas by the mean of the same and additionally done in a robust way by using the median absolute deviation and median, respectively. For all of the above features, missing values were dropped silently from the summary calculations. We were unable to use any of the columns produced by enabling the verboseColumns = TRUE option in findChromPeaks because all of the values returned were NAs.

We also calculated several novel metrics from the raw *m/z*/RT/intensity values by extracting the data points falling within each individual peak’s *m/z* and RT bounding box (values between the XCMS-reported min and max) separately for each file. The data points were then linearly scaled to fall within the 0-1 range by subtracting the minimum RT and dividing by the maximum RT, then each scaled RT was fit to a beta distribution with α values of 2.5, 3, 4, and 5, and a fixed β value of 5. This approach allowed us to approximate a bell curve with increasing degrees of right-skewness and the beta distribution was chosen because it is constrained between 0 and 1 and simple and speedy to generate in R. For each α value, Pearson’s correlation coefficient (*r*) was calculated between the beta distribution and the raw data, with the highest value returned as a metric for how peak-shaped the data were (Figure 8). The beta distribution with the highest *r* was also then used to estimate the noise level within the peak by scaling both the beta distribution probability densities and the raw data intensity values as described above, then subtracting the scaled beta distribution from the scaled intensity values, producing the residuals of the fit (Figure 8). The signal-to-noise ratio (SNR) was calculated by dividing the maximum original peak height by the standard deviation of the residuals multiplied by the maximum height of the original peak. This method of SNR calculation allowed us to rapidly estimate the noise within the peak itself rather than relying on background estimation using data points outside the peak, which may not exist or may be influenced by additional mass signals (Myers et al. 2017b). If there were fewer than 5 data points, a missing value was returned and dropped in subsequent summary calculations. Accessing the raw data values also allowed us to calculate the proportion of “missed” scans in a peak for which an RT exists at other masses in the same sample but for which no data was produced at the selected *m/z* ratio, divided by the total number of scans between the min and max RTs.

**Figure 8:**
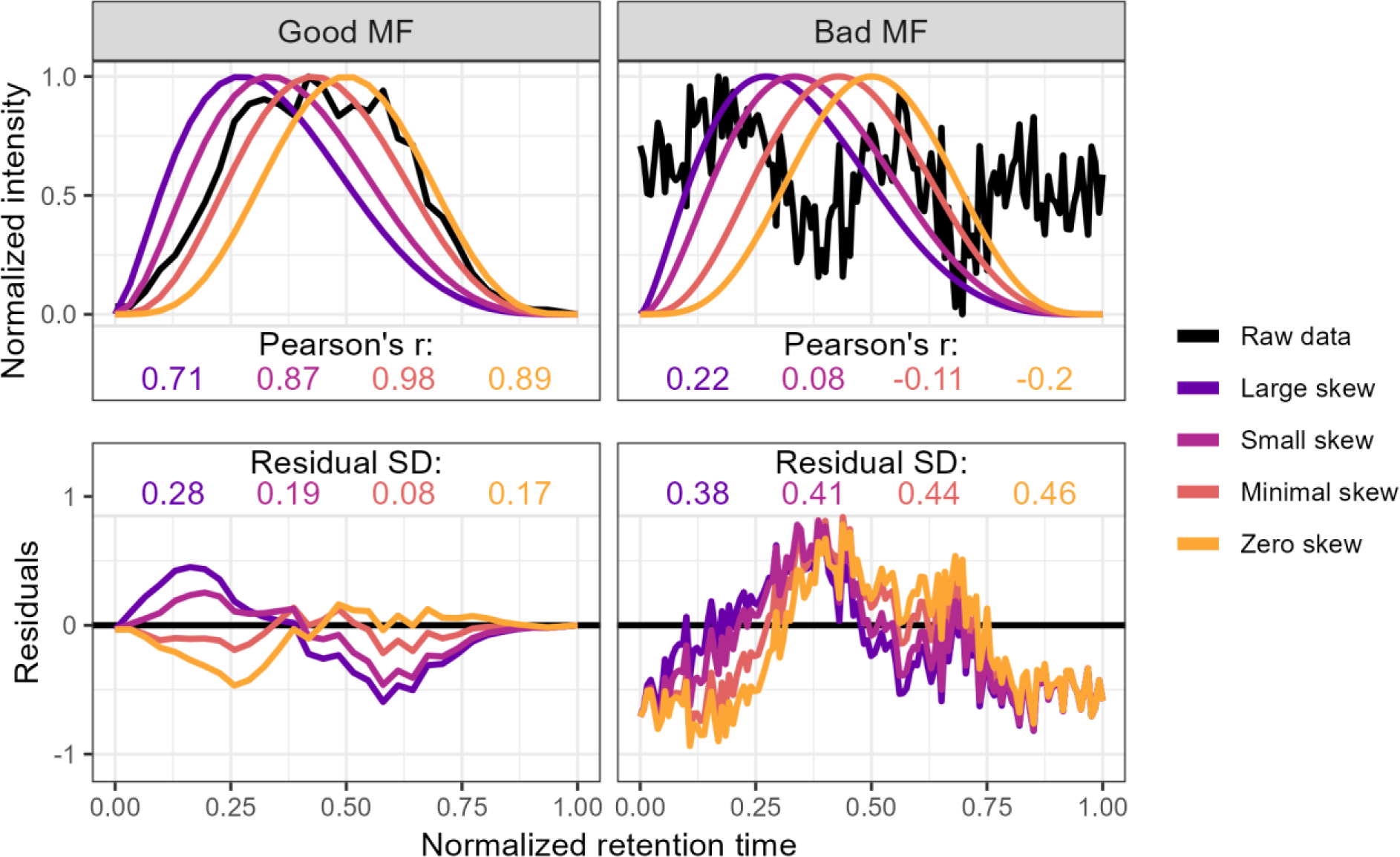
Method used to calculate the metrics for the two-parameter model from the raw data via comparison to an idealized pseudo-Gaussian peak for both manually identified “Good” and “Bad” peaks. Normalization was performed by linearly scaling the raw values into the 0-1 range by subtracting the minimum value and dividing by the maximum. Peak shape similarity was measured with Pearson’s correlation coefficient and the noise level is estimated as the standard deviation of the residuals after the raw data is subtracted from the idealized peak.

We additionally estimated the presence or absence of a ^13^C isotope using a similar method to extract the raw *m/z*/RT/intensity values within the peak bounding box, then searched the same RT values at an *m/z* delta of +1.003355 ± 4 PPM. In places where more than 5 data points existed at both the original mass and the ^13^C mass, we again used Pearson’s correlation coefficient to estimate the similarity between the two mass traces and used a trapezoidal Riemann sum to estimate the area of the original and isotope peaks. The overall feature isotope shape similarity was calculated by taking the median of the correlation coefficients. We also calculated the correlation coefficient of the ratio of the 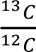peak areas across multiple files, expecting that a true isotope would have a fixed 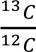ratio. Both the isotope shape similarity and the isotope area correlation were used as metrics in the downstream analysis. Peaks for which no isotope signal was detected or had too few scans to calculate the above metrics were imputed with NA values that were again dropped in the calculation of summary statistics for the mass feature as a whole. Because these isotope metrics typically had highly skewed distributions with most values very close to one, we normalized them by taking the log_10_ of one minus the value.

Distributions were visually inspected using a pairs plot and highly correlated (above a Pearson’s r ∼ 0.9) metrics had one of the redundant metrics removed.

### Regressions and model development

We used three different multiple logistic regression models to predict the likelihood of each MF being categorized as “Good”. The first model included all metrics calculated as described above in Methods, the second contained only those parameters immediately available from the XCMS output without revisiting the raw data (the four core peak metrics *m/z*, RT, peak width, area and their standard deviations plus the mysterious lmin, f, and scale values as well as the fraction of peaks, samples, and standards found), and the final model was a simple two-parameter model using only the peak shape and novel SNR metrics.

In each case, we categorized each mass feature as a true positive (TP) if it was predicted to be Good and was manually classified as Good, a true negative if both predicted and classified as Bad, a false positive if predicted to be Good but manually classified as Bad, and a false negative if predicted to be Bad but was in fact manually classified as Good. This allowed us to additionally define two useful measures of success, the traditionally-defined false discovery rate (FDR, defined as 1-precision or the number of false positives divided by the total number of predicted positives) and the percentage of good features found (GFF, also known as the recall or sensitivity and defined as the number of true positives divided by the total number of actual positives).

To further explore questions of model stability and the potential for overfitting, we compared the predictions from a Falkor-trained model to a MESOSCOPE-trained model. This comparison was done in both the raw probability space as well as a rank-ordered space to test whether the most extreme likelihood (i.e. very best and very worst) MFs were consistently found to be most extreme independently of the actual likelihood predicted. For the raw probability space we compared the predictions using Pearson’s correlation coefficient, while Spearman’s rank-ordered coefficient was used for the ranked space. We additionally looked at the estimates produced by these two models and compared them with the combined model trained on both datasets combined to assess the model stability directly.

We also measured the robustness of the model under a smaller training set, emulating a situation in which only a fraction of the data was available or only a portion of the mass features had been labeled. This allowed us to test the required sample size for the different models, with a larger sample size presumably required for the models with more parameters. Because no parameter was present in all 3 models, we looked at the top 2 most significant parameters from each model: average *m/z* and peak shape for the full model, average *m/z* and the standard deviation in retention time for the XCMS model, and peak shape and SNR for the two-parameter model.

Finally, we tested whether the performance could be improved with regularized regression or random forest models. These models handle correlated variables better than ordinary least squares regression, so we also included several additional implementations of the peak shape and novel SNR parameters when summarizing across multiple files, using a max and a median of the top-three best values rather than just the overall median as well as a log-transformed version of the median peak shape calculated as *median*(*log*_10_(1 − *r*)) where *r* is Pearson’s correlation coefficient, as described above (Figure 2). Cross-validation was used to select the optimal tuning parameter λ with glmnet package’s cv.glmnet for an elastic net penalty (α) of 0, 0.5, and 1. Random forests were implemented using the randomForest package with default settings and a factor-type response vector to ensure classification was applied rather than regression.

### Application of the model to novel datasets

After exploring the different models described above and determining that the two-parameter model would likely perform most consistently on novel datasets, we applied this trained model on two additional datasets that differed significantly from the training data. The CultureData dataset was produced in the Ingalls lab like MESOSCOPE and Falkor, but represent data from a variety of phytoplankton and bacterial cultures in fresh and salt water rather than environmental samples.

The Pttime dataset was discovered on Metabolomics Workbench where it has been assigned Project ID PR001317. The data can be accessed directly via it’s Project DOI: 10.21228/M8GH6P. This project dataset consists of *Phaeodactylum tricornutum* cultures collected at a variety of timepoints from both pelleted cells and the released exudate. This dataset was chosen because of the similar LC-MS setup used as a benchmark for the performance that other labs with similar setups may expect to achieve using the trained model directly.

Each of these datasets were only fractionally labeled, with those MFs above the 0.9 likelihood threshold according to the two-parameter model reviewed manually and categorized. This stricter threshold was chosen because we felt less comfortable interpreting results based on mass features that were only 50% likely to be real, but did not feel the need to be so strict with this exploratory analysis that we wanted to limit it to 99+% likelihood MFs.

### Using variable thresholds to determine effects on biological conclusions

We explored the implications of applying this model to the MESOSCOPE dataset at a variety of thresholds. In univariate space, we used nonparametric Kruskal-Wallis analyses of variance to measure the difference between the surface (15m), DCM (∼110m), and 175m samples because the metabolite peak areas could not be assumed to be normally distributed. These univariate tests were then controlled for multiply hypothesis testing using R’s p.adjust function with method fdr (Benjamini and Hochberg 1995). We also performed post-hoc Dunn tests provided by the rstatix package to categorize the response to depth for those mass features for which the KW test was significant, with responses falling into one of the 14 classes possible when permuting the sign and significance of the Dunn test outputs (Dunn 1964). p-values obtained from the Dunn tests were not FDR controlled because it was used as a categorization tool rather than a null hypothesis test. In multivariate space, we used a permutational MANOVA (PERMANOVA) (Anderson 2017) provided by the vegan package’s adonis2 function to test for multivariate differences in structure of the metabolome with depth (Oksanen et al. 2022). We ran multiple PERMANOVAs with a different subset of mass features included each time, corresponding to using the output from XCMS directly, likelihood thresholds of 0.01, 0.1, 0.5, 0.9, and finally only those MFs manually annotated as good.

All analyses were run in R (R Core Team 2022), version 4.2.2, and code is available on GitHub at https://github.com/wkumler/MS_metrics.

## Declarations

### Ethics approval and consent to participate

Not applicable

### Consent for publication

Not applicable

## Availability of data and materials

The raw mzML files are all available on Metabolomics Workbench. The Falkor and MESOSCOPE datasets can be found under project ID PR001738 via http://dx.doi.org/10.21228/M82719. The CultureData samples were appended to the previously existing culturing collection, accessible at project ID PR001021 via http://dx.doi.org/10.21228/M8QM5H. Pttime is located under Project ID PR001317 and can also be accessed directly using its Project DOI: http://dx.doi.org/10.21228/M8GH6P. Code and other raw data is available on the GitHub repository at https://github.com/wkumler/MS_metrics. The manuscript has been rendered as a single R Markdown document with analyses contained within for reproducibility.

## Authors’ contributions

WK extracted the Falkor samples, processed the data, performed the analyses, and wrote the manuscript. BJH helped design the metrics and implement the regressions as well as providing support and context for the analysis. AEI provided funding and data and helped to interpret the conclusions and edit the manuscript.

## Acknowledgements

This work was supported by the University of Washington eScience Institute through their Data Science Incubator program. The authors would also like to acknowledge Laura Carlson for her expertise in obtaining the CultureData, MESOSCOPE, and Falkor datasets; Katherine Heal, Angie Boysen, and Bryn Durham for their assistance with culturing and collecting the cultured samples; the captain and crew of the R/Vs *Kilo Moana* and *Falkor*, Wei Qin, Rachel Lundeen, and the SCOPE ops team for collecting and processing the environmental samples; Joshua Sacks, Dave Beck, and the entire eScience incubator team for their feedback on the project development and scope; and Brisson Vanessa and LLNL for making their Pttime dataset available for reuse on Metabolomics Workbench. Additionally, the R packages plotly, dbscan, ClusterR, RaMS, and the entire tidyverse were crucial for preliminary data exploration.

## Supplement

Supplemental figure 1: Distribution and single-parameter logistic curves for each metric extracted for model training, shown separately for the MESOSCOPE and Falkor datasets. Histograms show the distribution of good and bad mass features by color across the span of the data on the x-axis with the number of MFs in each bin shown on the y-axis. Scatterplots show the same x-axis but show the results of a logistic regression on the single parameter, with the line of best fit in black and a ±1 standard error ribbon around it in grey. Vertical jittering has been applied when plotting to reduce the number of overlapping points.

Supplemental figure 2: Model parameter estimates for each of the metrics in the full model, additionally broken down by their inclusion in the two-parameter (raw_data) and XCMS-exclusively models. Colors correspond to the dataset used to train the logistic regression model, with “both” indicating a combined model using all manually-labeled features across both datasets.

Supplemental figure 3: Robustness of the two most significant metrics across the full (all_params), XCMS-only (xcms_params), and two-parameter (raw_data_params) models. The x-axis corresponds to the fraction of the data used to train the model and the y-coordinate shows the estimated value for the specified term in the subset across 10-fold replicated subsampling. The grey bar in the background corresponds to the estimate of the full model +/-1SE (thinner dark grey bar) and 2SE (thicker light grey bar).

Supplemental figure 4: Performance of regularized regression and random forest models on internally (same train-test) and externally (different train-test) validated datasets.

## Abbreviations

DCM: Deep Chlorophyll Maximum
EIC: Extracted Ion Chromatogram
FDR: False Discovery Rate
GFF: Good Feature Found
HILIC: Hydrophilic Interaction Liquid Chromatography
LC: Liquid Chromatography
MF: Mass Feature
MS: Mass Spectrometry
PPM: parts-per-million
RT: Retention time
SNR: Signal to Noise Ratio

