## Supplemental Figure 1 for "Picky with peakpicking: assessing chromatographic peak quality with simple metrics in metabolomics"

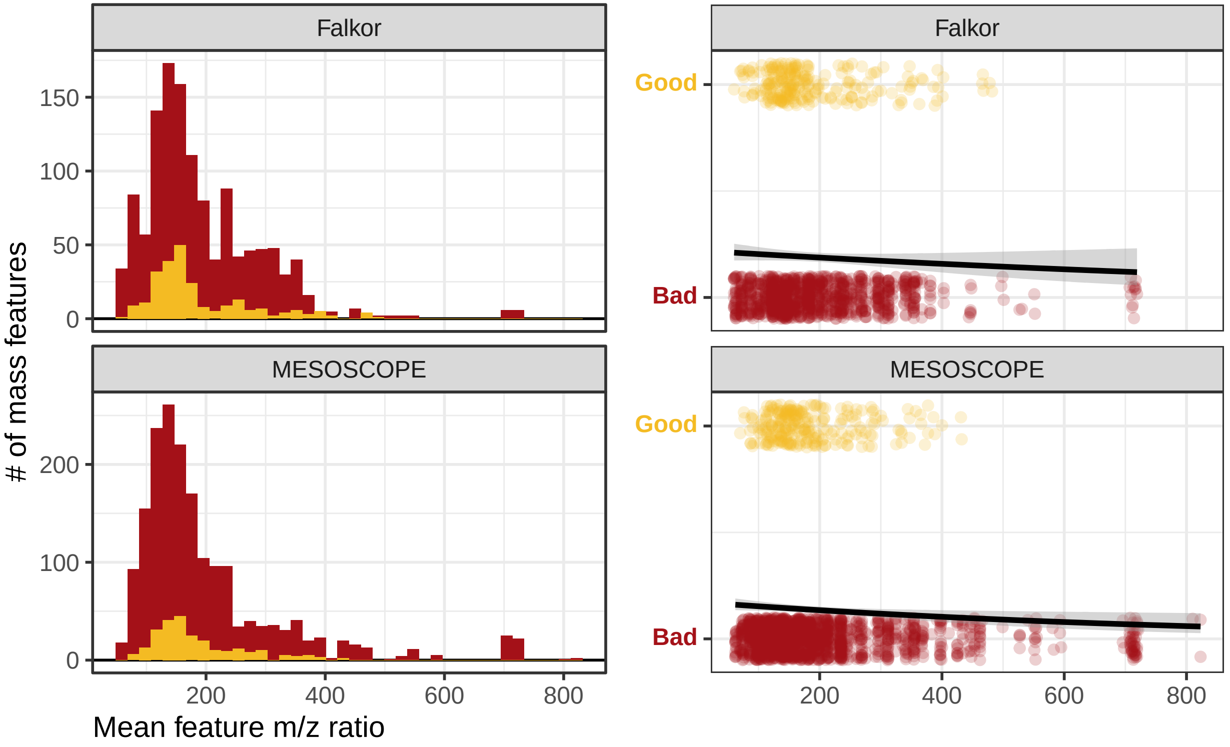

Falkor
MESOSCOPE
Falkor
MESOSCOPE
150
100
50
0
200
100
0
Good
Bad
Good
Bad
200
400
600
800
### of mass features
Mean feature m/z ratio

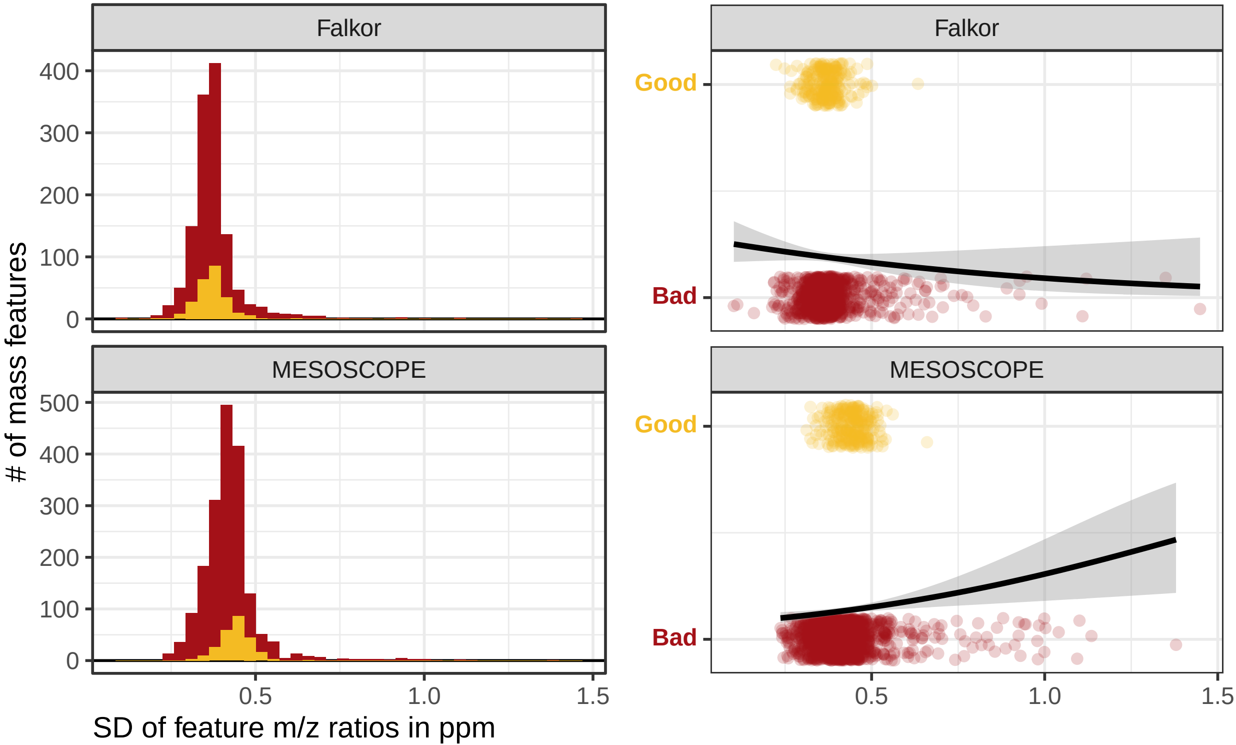
Falkor
MESOSCOPE
Falkor
MESOSCOPE
Good
Bad
Good
Bad
### of mass features
400
300
200
100
0
500
400
300
200
100
0
0.5
1.0
1.5
0.5
1.0
1.5
SD of feature m/z ratios in ppm

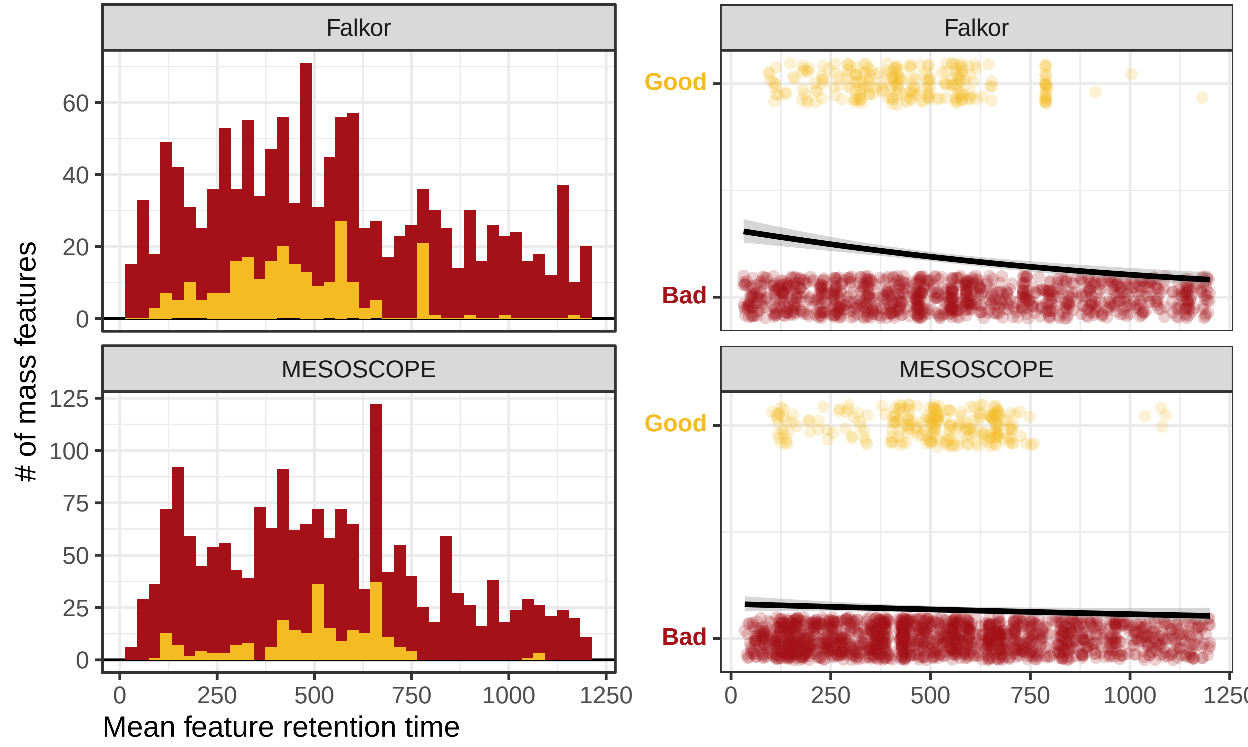

Falkor
MESOSCOPE
### of mass features
Mean feature retention time
Good
Bad
0
20
40
60
25
50
75
100
125
250
500
750
1000
1250

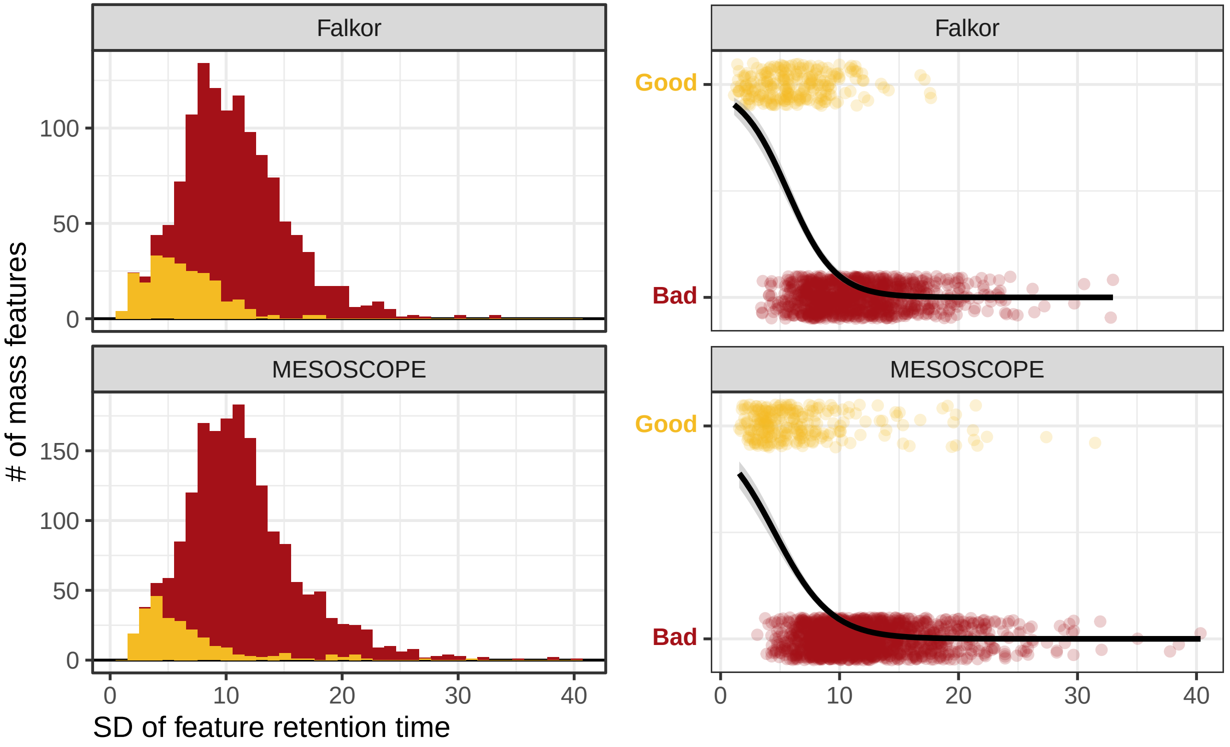

Falkor
MESOSCOPE
Falkor
MESOSCOPE
### of mass features
SD of feature retention time
Good
Bad
Good
Bad
0
50
100
150
0
10
20
30
40

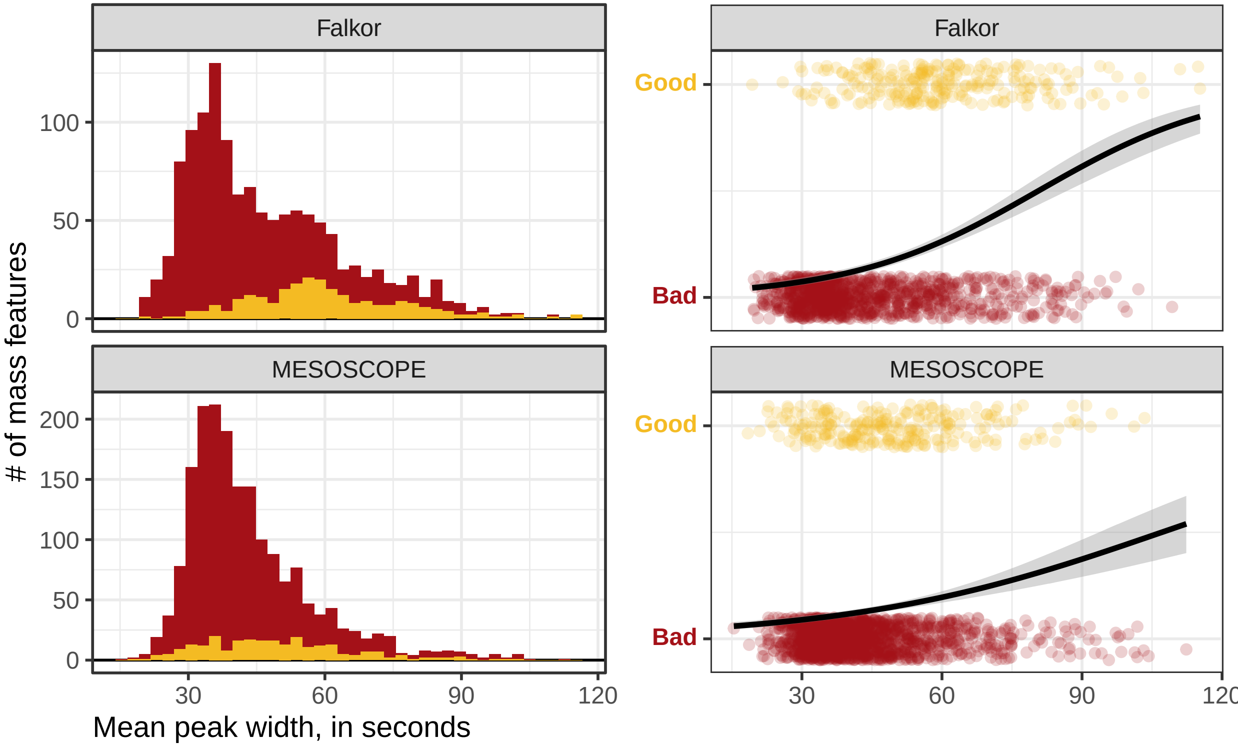

Falkor
MESOSCOPE
Falkor
MESOSCOPE
Good
Bad
Good
Bad
0
50
100
0
50
100
150
200
30
60
90
120
### of mass features
Mean peak width, in seconds

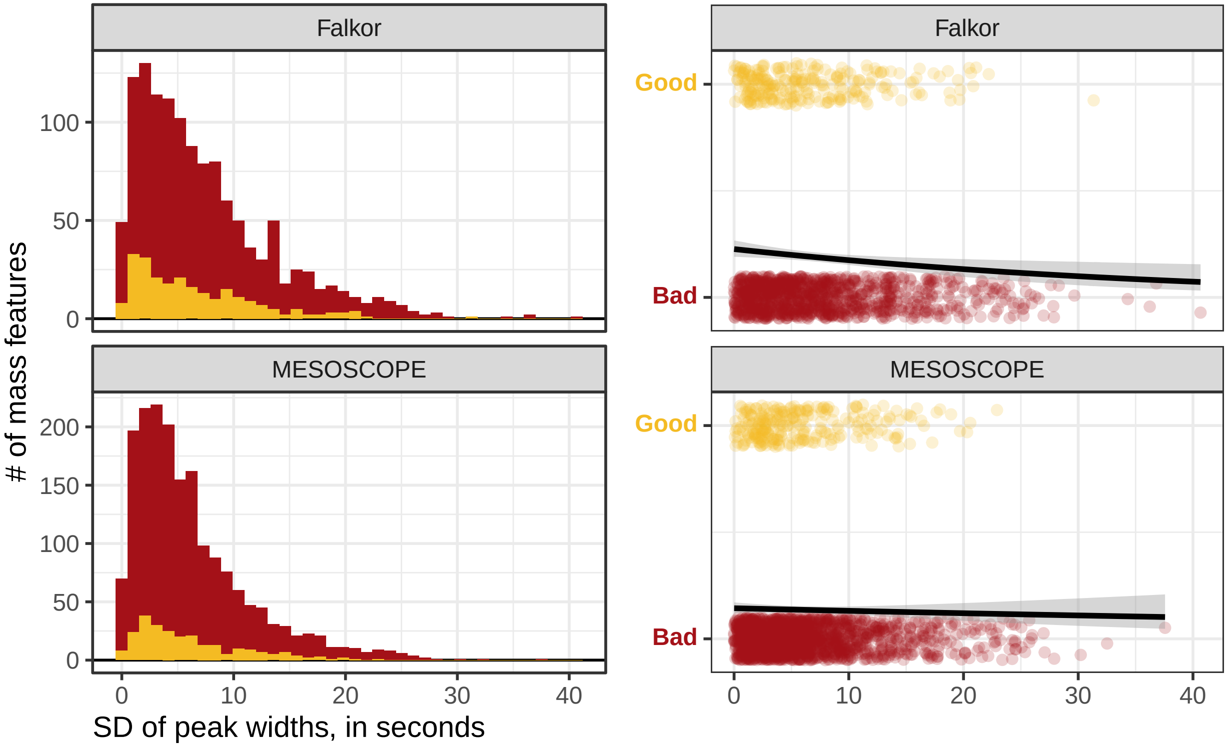
Falkor
MESOSCOPE
### of mass features
100
50
0
200
150
100
50
0
0
10
20
30
40
SD of peak widths, in seconds
Good
Bad
Falkor
Good
Bad
MESOSCOPE
0
10
20
30
40

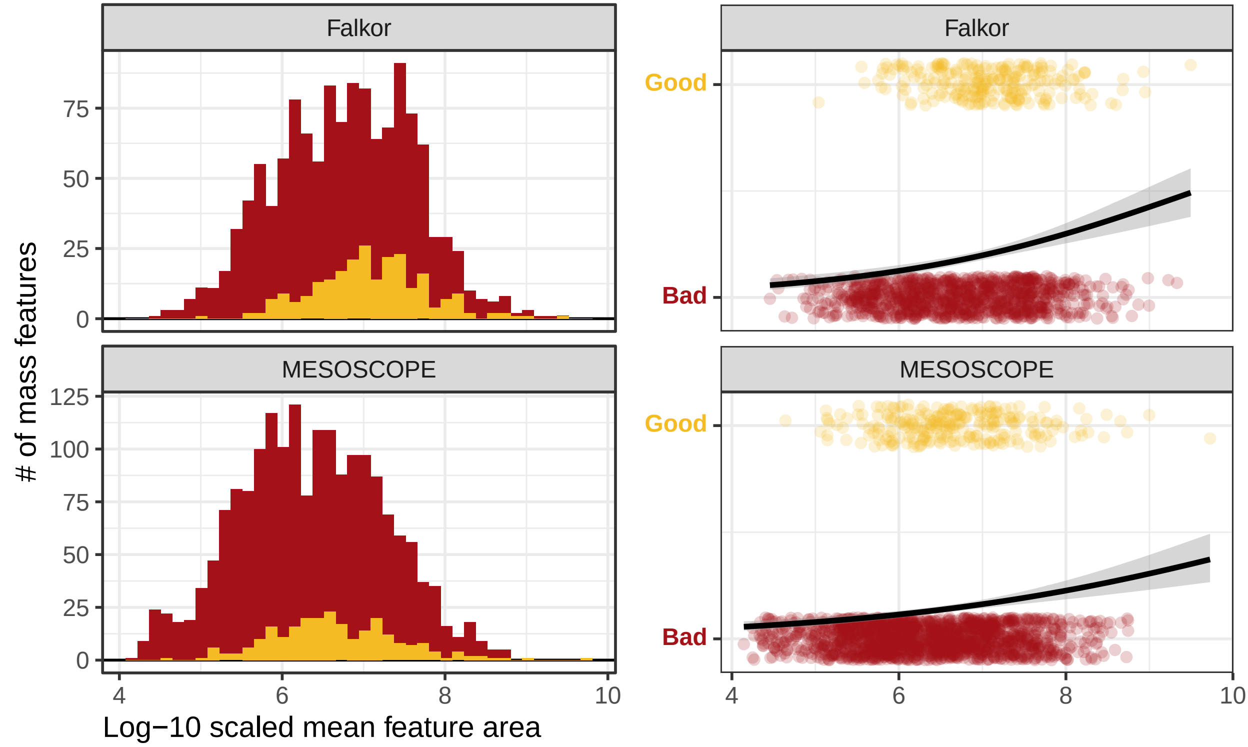

Falkor
MESOSCOPE
Falkor
MESOSCOPE
### of mass features
Log−10 scaled mean feature area
Good
Bad
Good
Bad
0
25
50
75
0
25
50
75
100
125
4
6
8
10
4
6
8
10

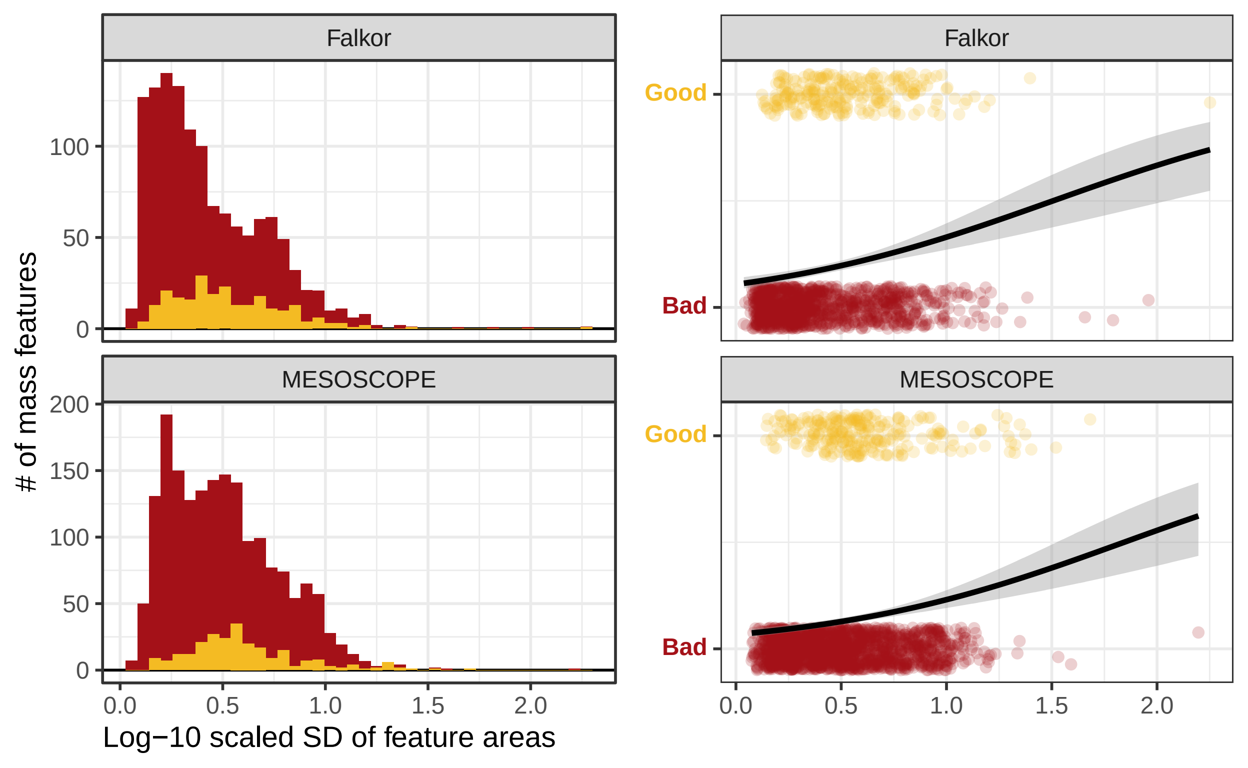

Falkor
MESOSCOPE
Falkor
MESOSCOPE
### of mass features
0
50
100
150
200
0.0
0.5
1.0
1.5
2.0
Log−10 scaled SD of feature areas
Good
Bad

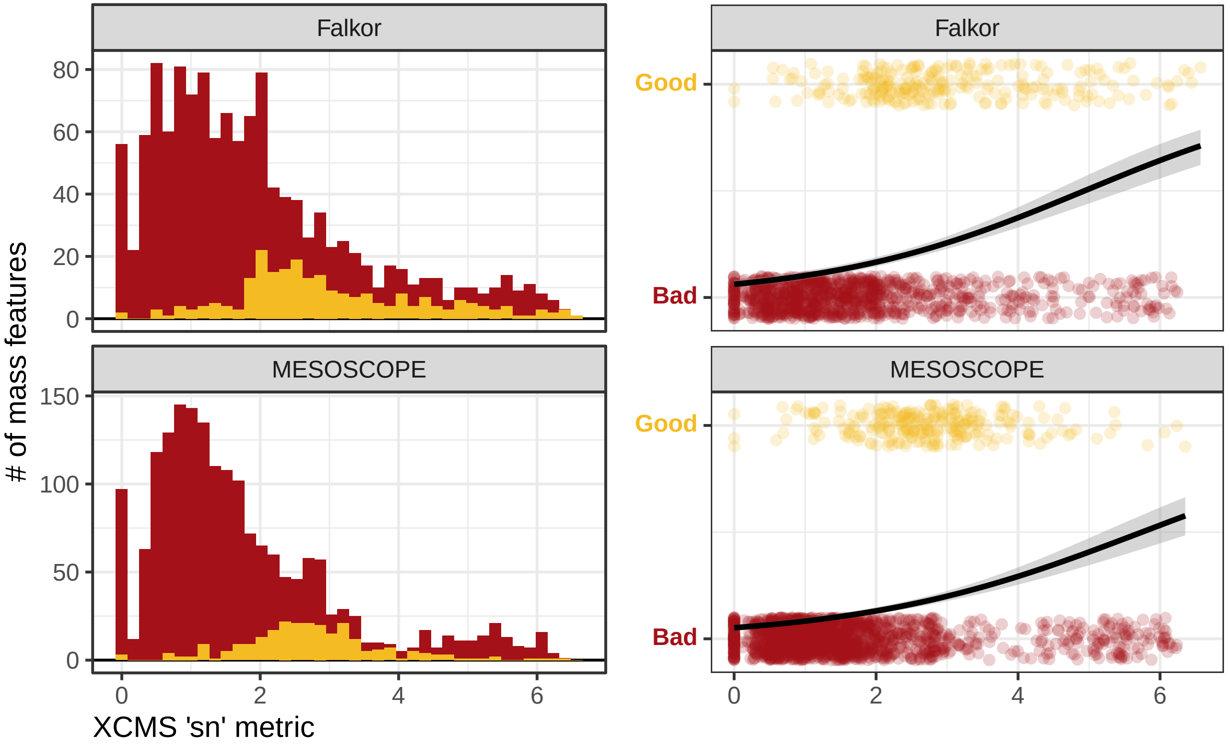

Falkor
MESOSCOPE
### of mass features
XCMS 'sn' metric
Good
Bad
0
20
40
60
80
50
100
150
2
4
6

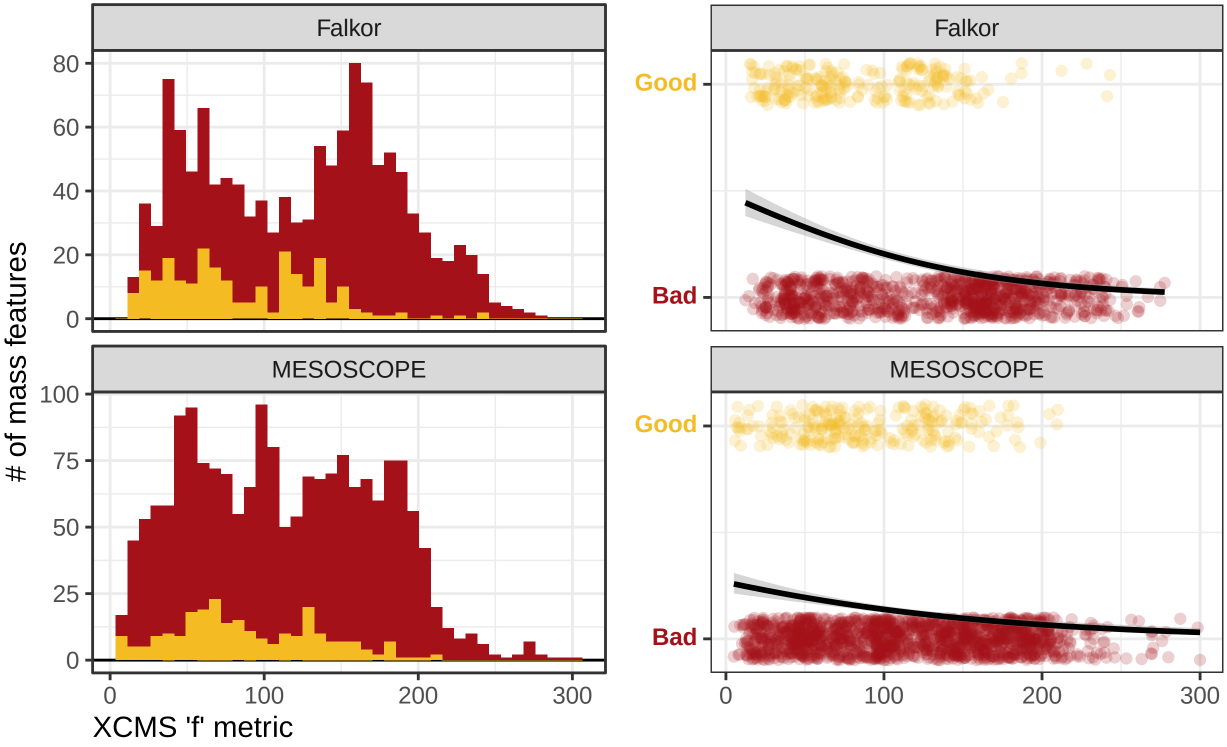

Falkor
MESOSCOPE
Falkor
MESOSCOPE
### of mass features
XCMS 'f' metric
Good
Bad
Good
Bad
80
60
40
20
0
100
75
50
25
0
0
100
200
300
0
100
200
300

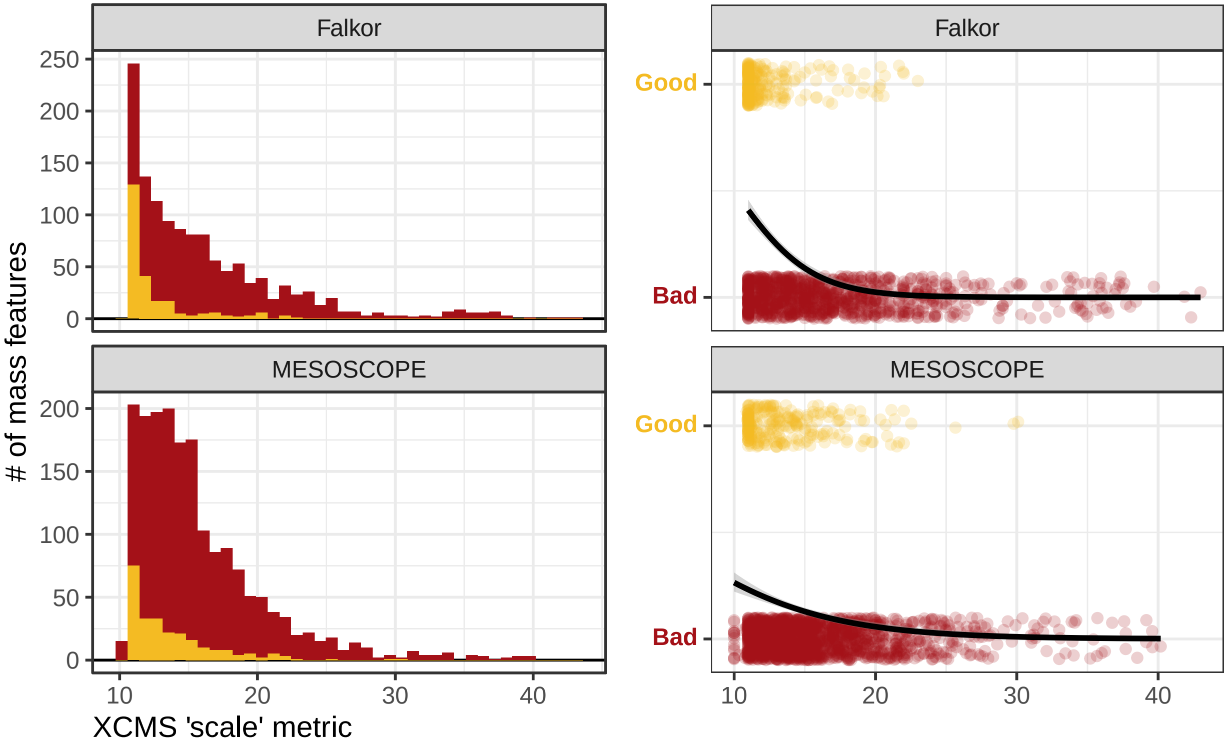
Falkor
MESOSCOPE
### of mass features
XCMS 'scale' metric
Good
Bad
0
50
100
150
200
250
10
20
30
40

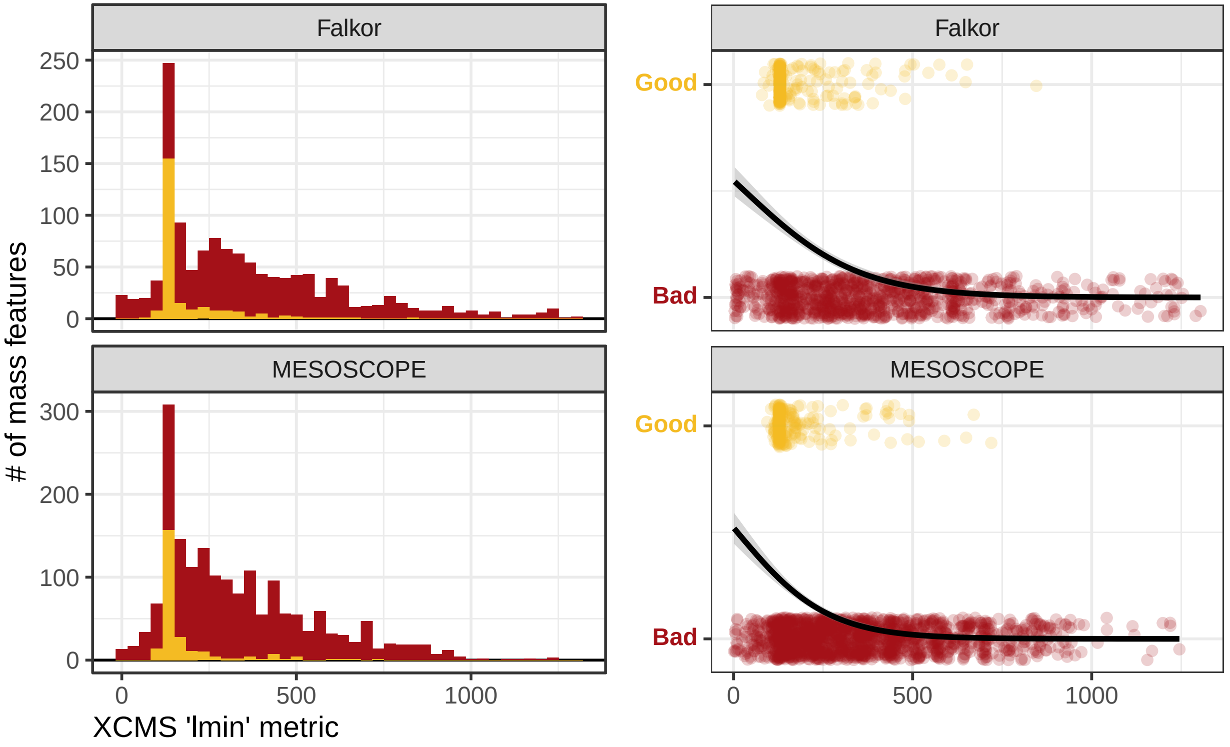
Falkor
MESOSCOPE
### of mass features
XCMS 'lmin' metric
0
50
100
150
200
250
300
500
1000
Good
Bad

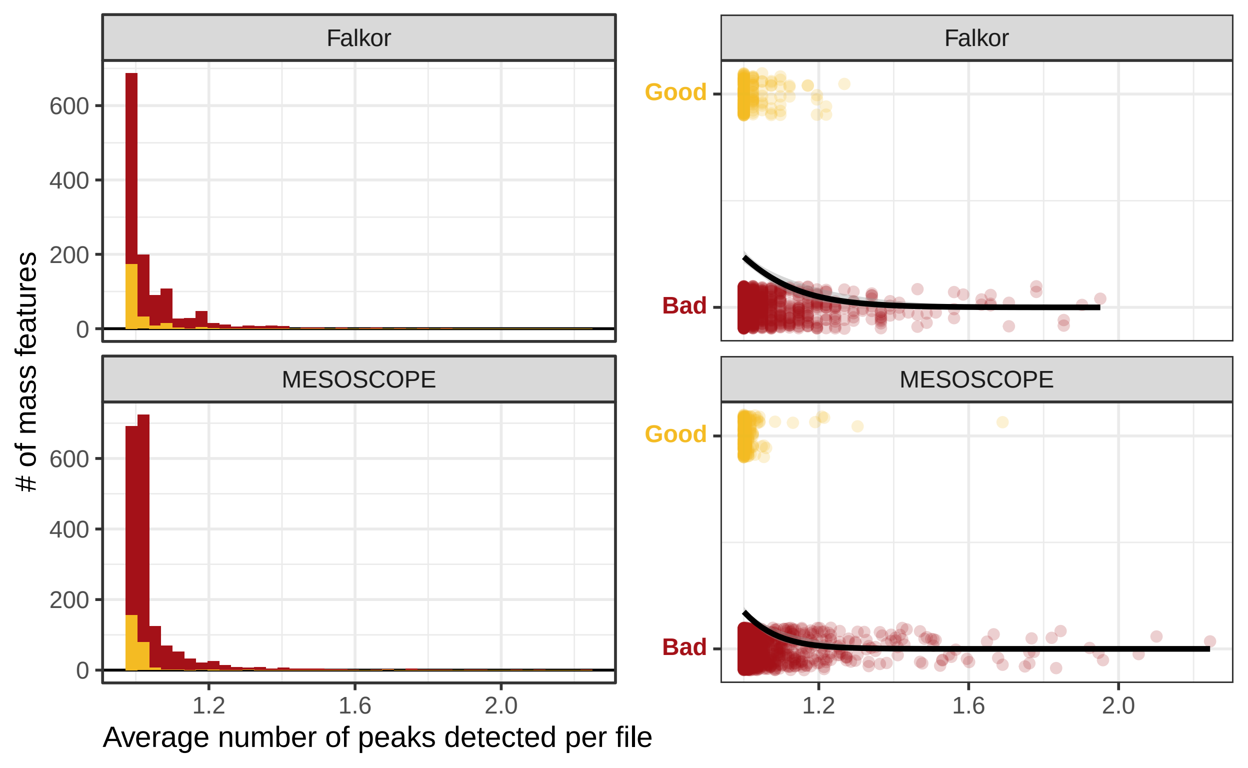
Falkor
MESOSCOPE
Good
Bad
600
400
200
0
1.2
1.6
2.0
### of mass features
Average number of peaks detected per file

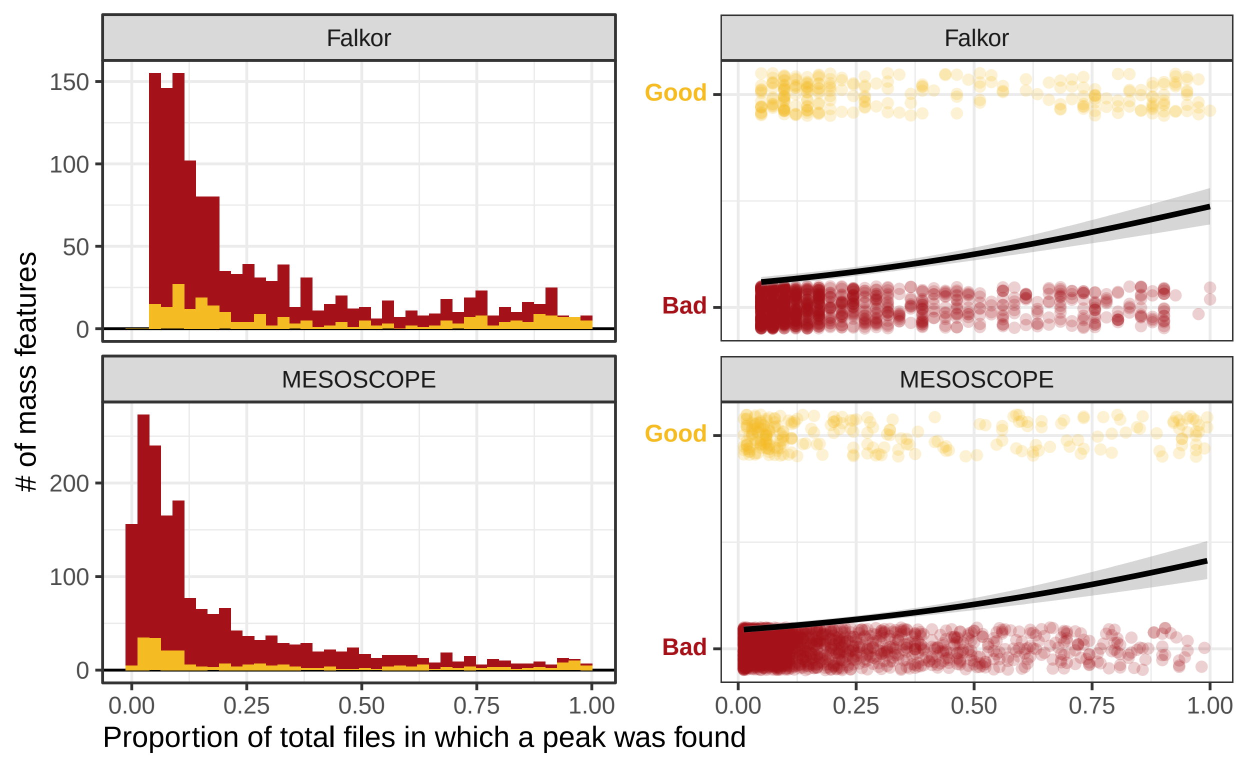
Falkor
MESOSCOPE
Falkor
MESOSCOPE
Good
Bad
Good
Bad
### of mass features
150
100
50
0
200
100
0
0.00
0.25
0.50
0.75
1.00
0.00
0.25
0.50
0.75
1.00
Proportion of total files in which a peak was found

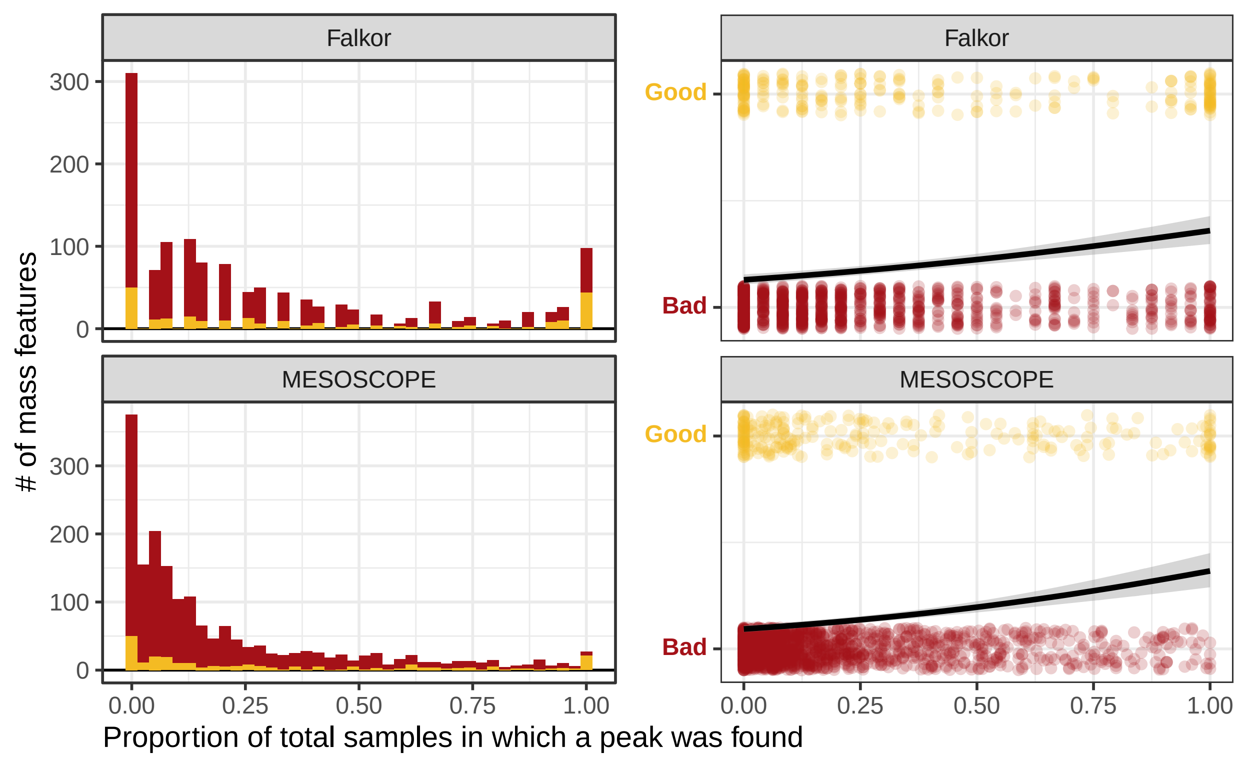

Falkor
Falkor
MESOSCOPE
MESOSCOPE
300
200
100
0
300
200
100
0
Good
Bad
Good
Bad
### of mass features
0.00
0.25
0.50
0.75
1.00
0.00
0.25
0.50
0.75
1.00
Proportion of total samples in which a peak was found

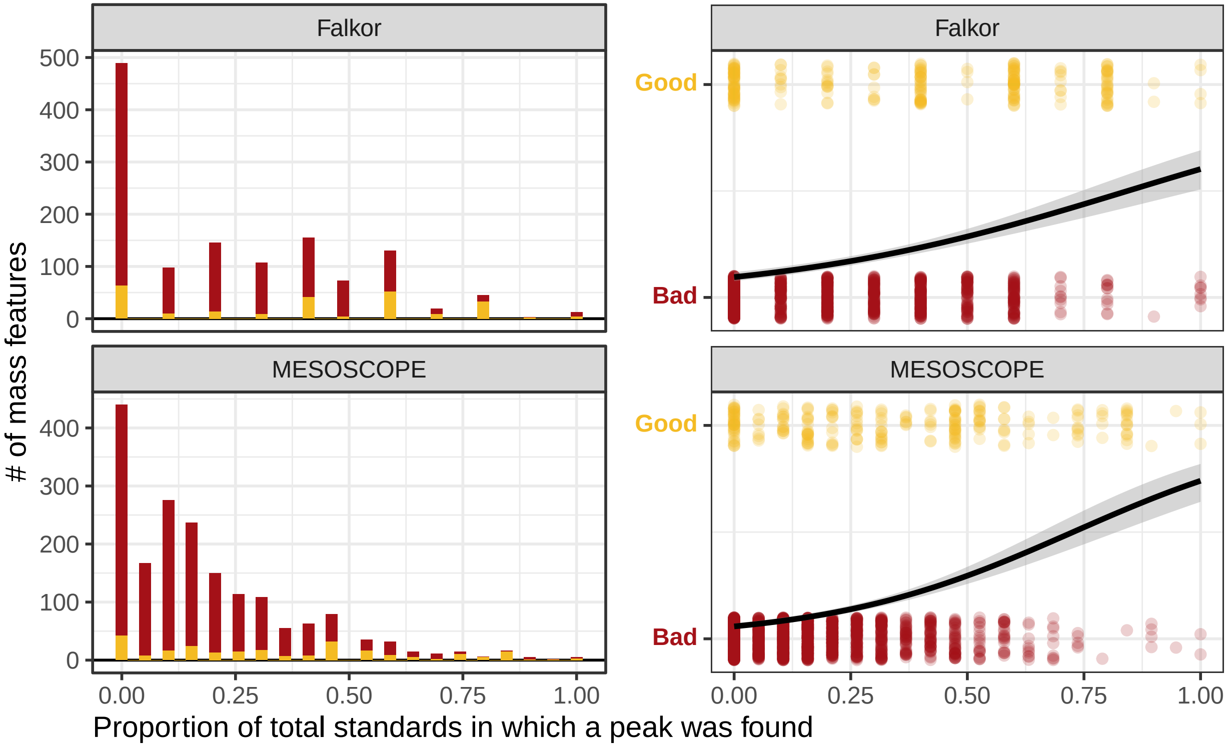

Falkor
MESOSCOPE
Good
Bad
0
100
200
300
400
500
0.00
0.25
0.50
0.75
1.00
### of mass features
Proportion of total standards in which a peak was found

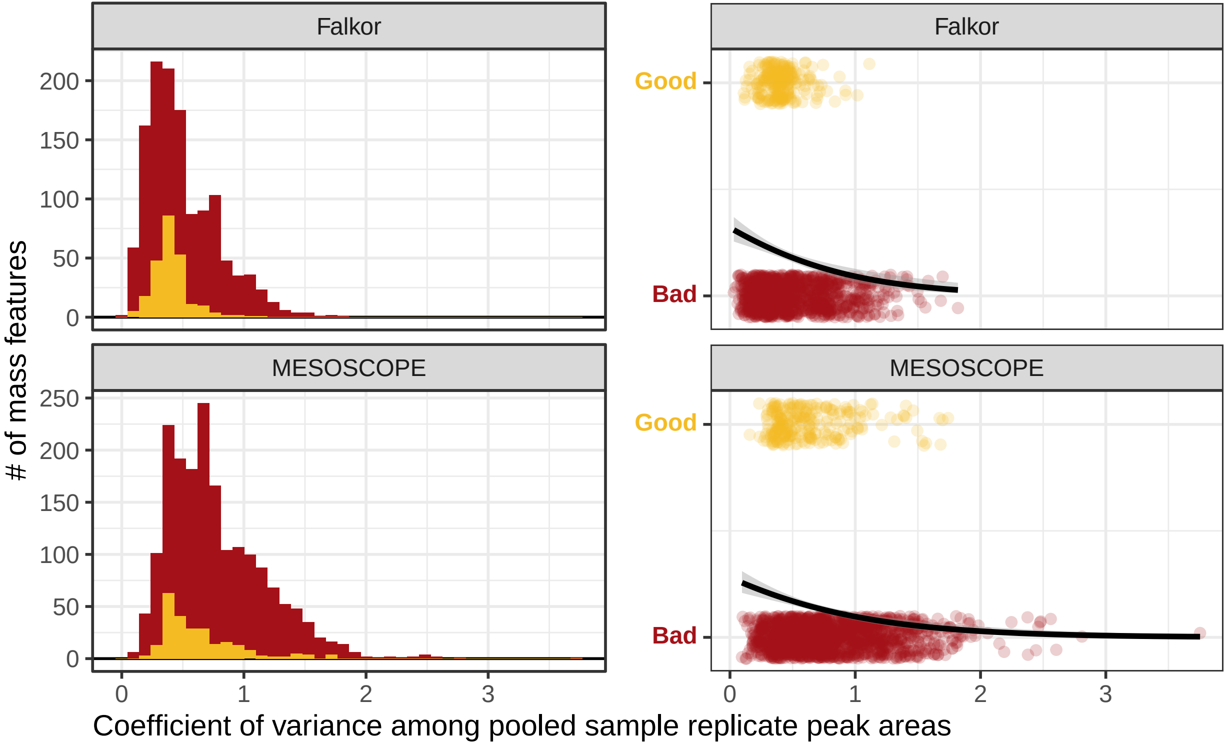
Falkor
Falkor
MESOSCOPE
MESOSCOPE
### of mass features
200
150
100
50
0
250
200
150
100
50
0
Good
Bad
Good
Bad
0
1
2
3
0
1
2
3
Coefficient of variance among pooled sample replicate peak areas

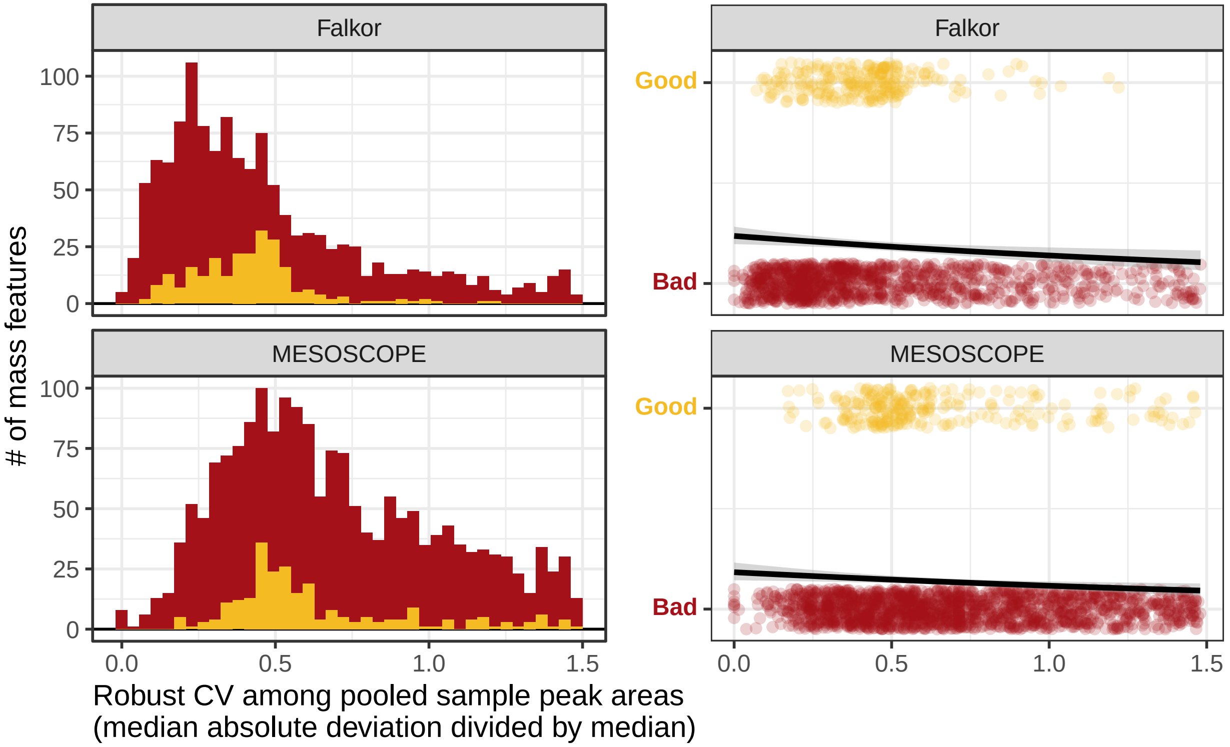

Falkor
MESOSCOPE
Falkor
MESOSCOPE
100
75
50
25
0
0.0
0.5
1.0
1.5
Good
Bad
### of mass features
Robust CV among pooled sample peak areas
(median absolute deviation divided by median)

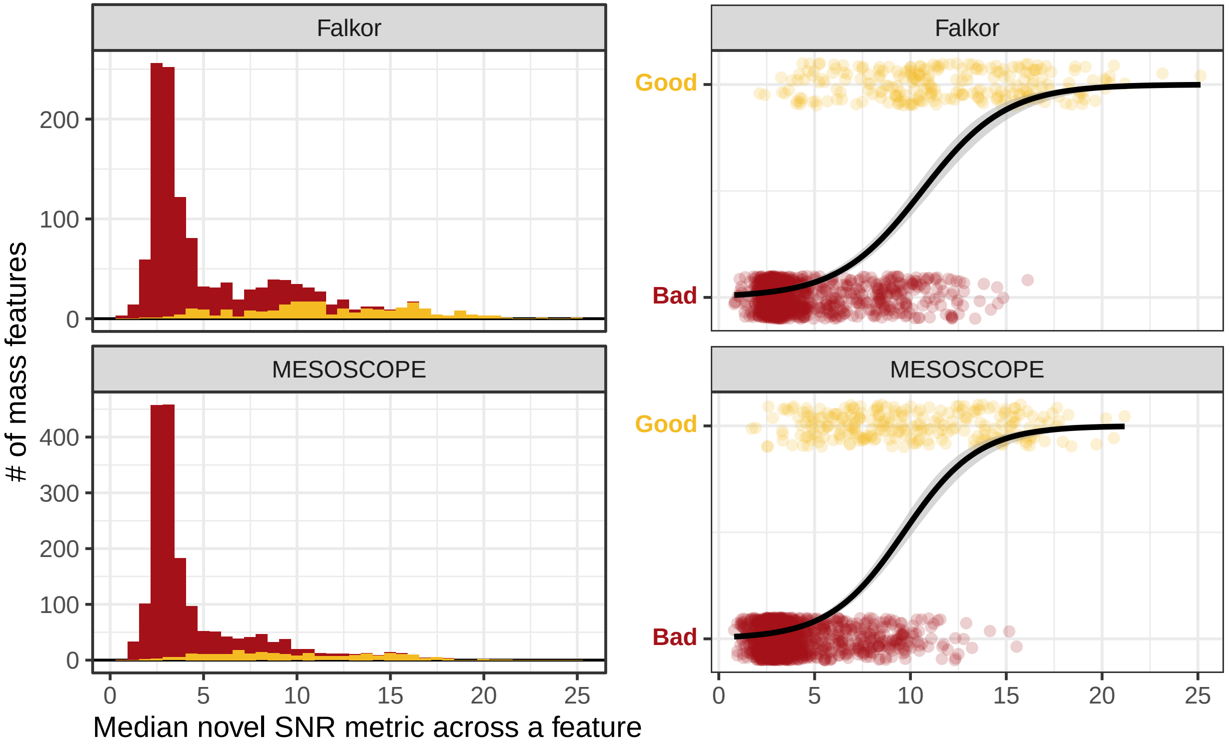

Falkor
MESOSCOPE
Falkor
MESOSCOPE
### of mass features
Median novel SNR metric across a feature
Good
Bad
Good
Bad
0
100
200
300
400
0
5
10
15
20
25

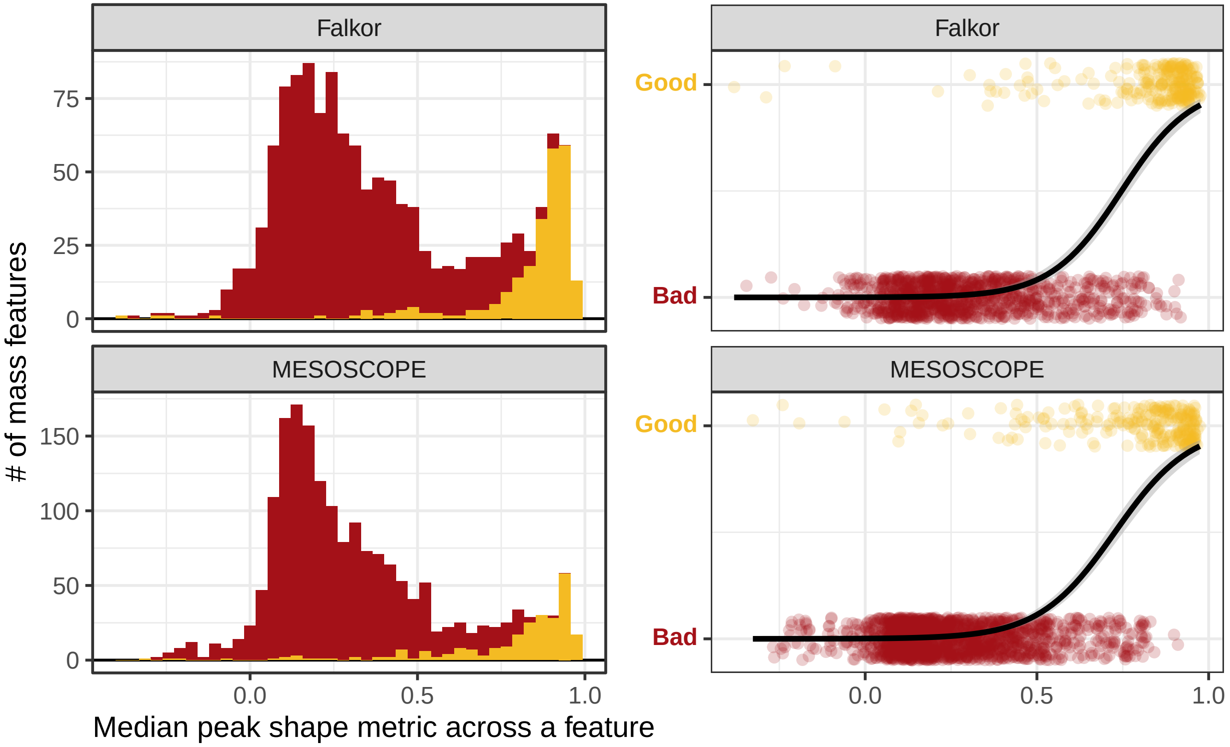

Falkor
MESOSCOPE
Falkor
MESOSCOPE
Good
Bad
Good
Bad
75
50
25
0
150
100
50
0
0.0
0.5
1.0
0.0
0.5
1.0
### of mass features
Median peak shape metric across a feature

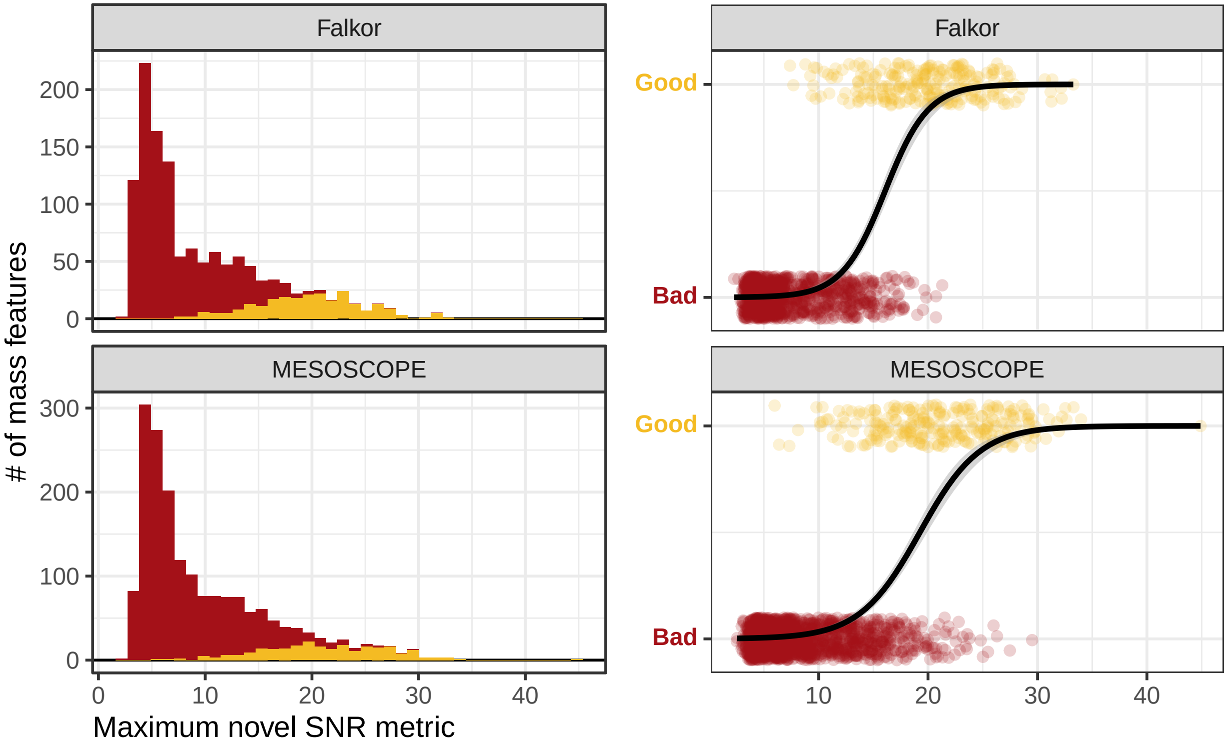

Falkor
MESOSCOPE
Falkor
MESOSCOPE
### of mass features
200
150
100
50
0
300
200
100
0
0
10
20
30
40
Maximum novel SNR metric
Good
Bad
Good
Bad
10
20
30
40

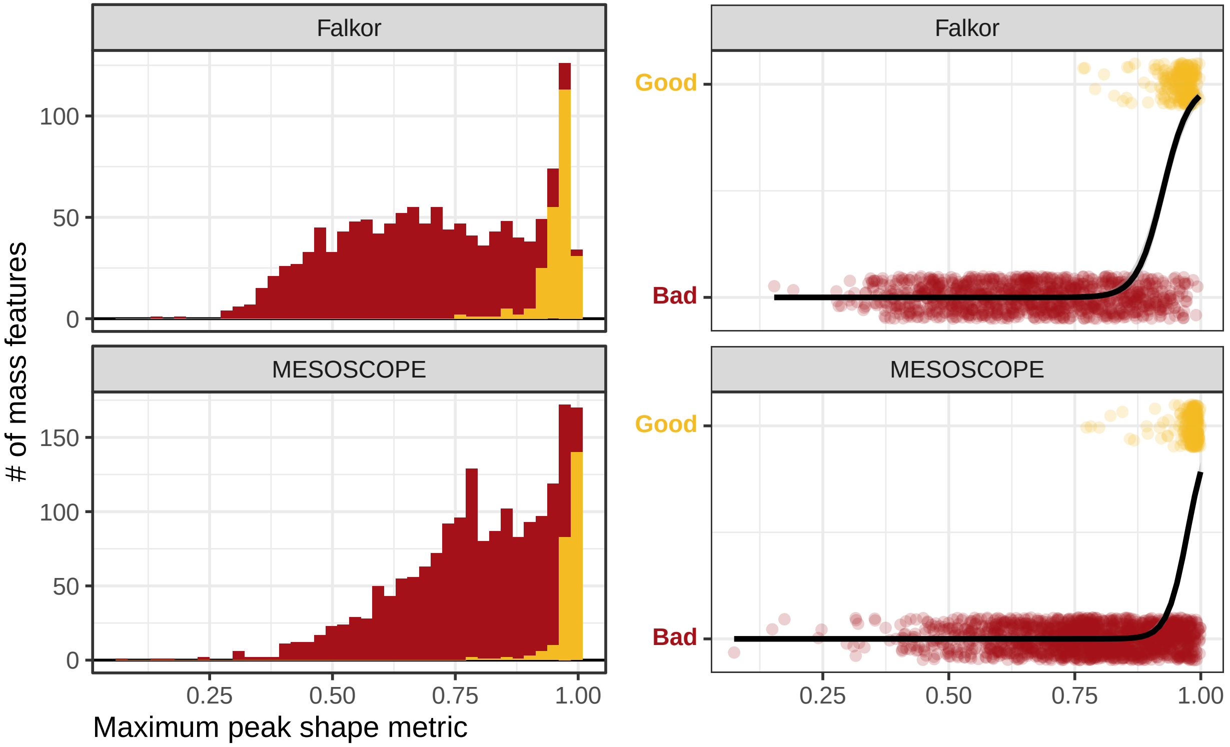
Falkor
MESOSCOPE
Falkor
MESOSCOPE
### of mass features
Maximum peak shape metric
Good
Bad
Good
Bad
0
50
100
0
50
100
150
0.25
0.50
0.75
1.00
0.25
0.50
0.75
1.00

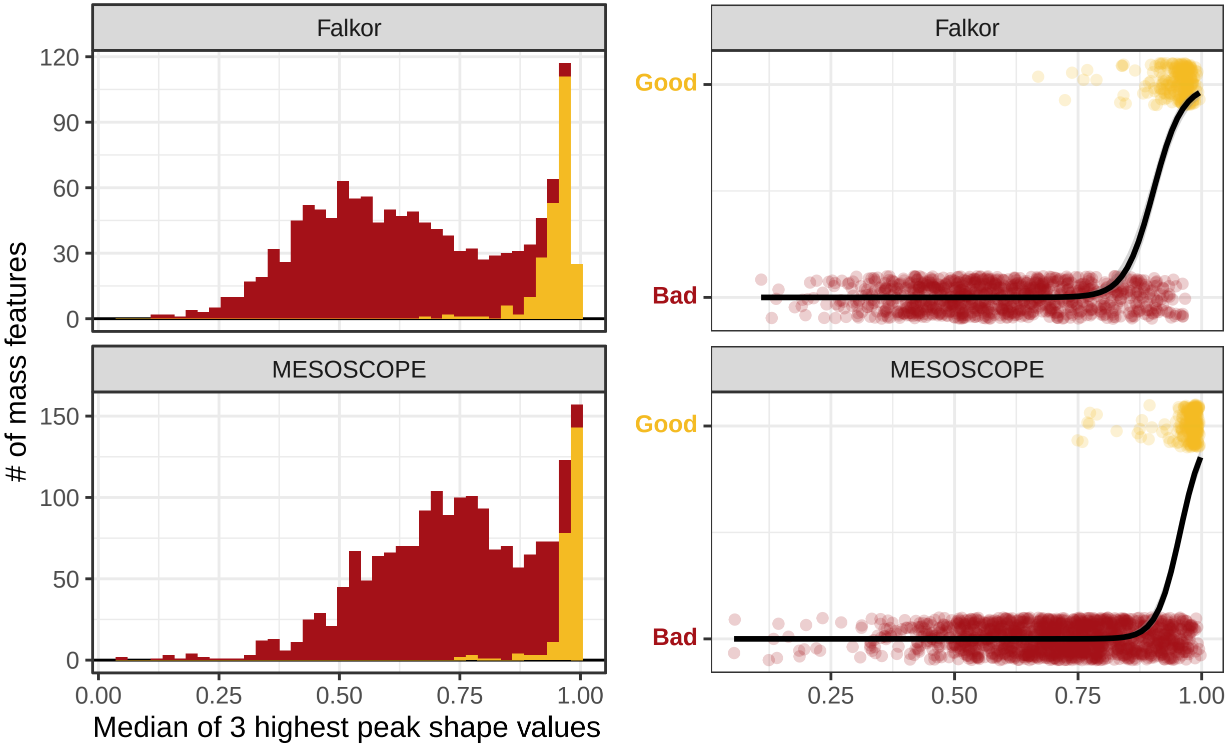

Falkor
MESOSCOPE
### of mass features
Median of 3 highest peak shape values
120
90
60
30
0
150
100
50
0
0.00
0.25
0.50
0.75
1.00
Falkor
MESOSCOPE
Good
Bad
Good
Bad
0.25
0.50
0.75
1.00

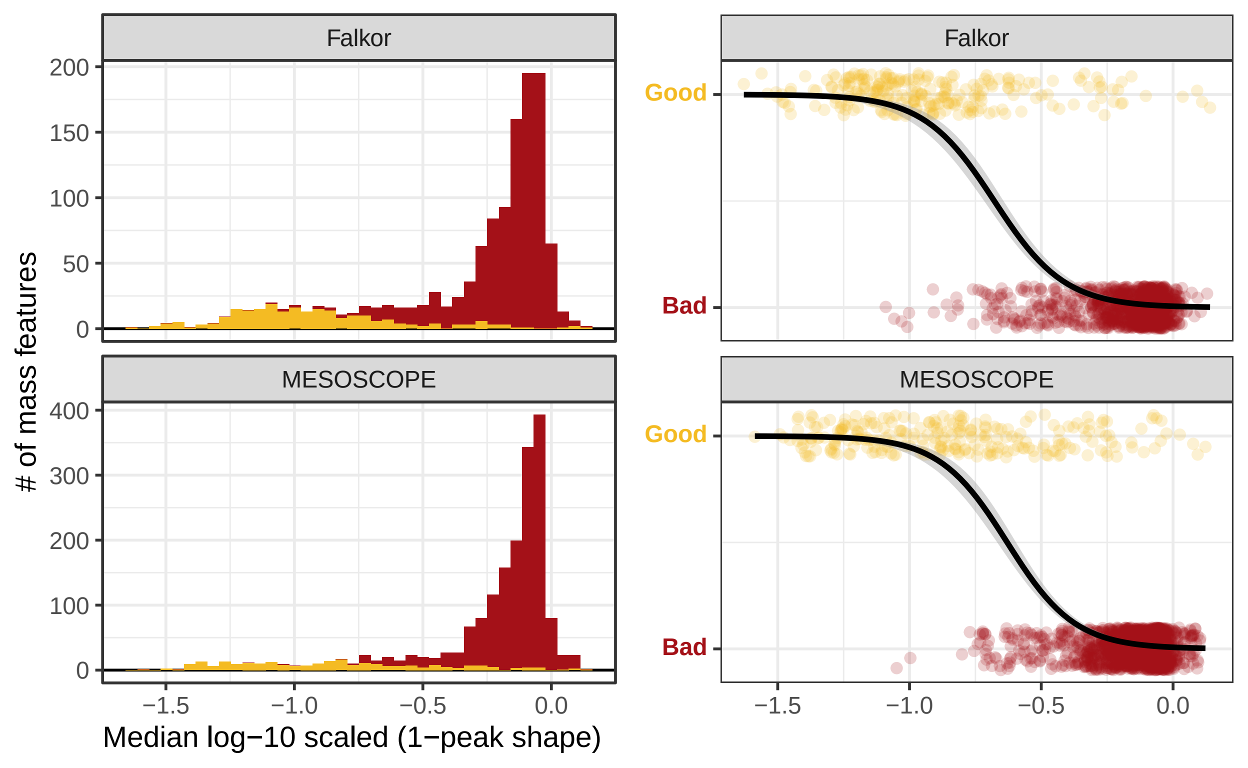
Falkor
MESOSCOPE
Good
Bad
### of mass features
Median log−10 scaled (1−peak shape)
200
150
100
50
0
400
300
200
100
0
−1.5
−1.0
−0.5
0.0

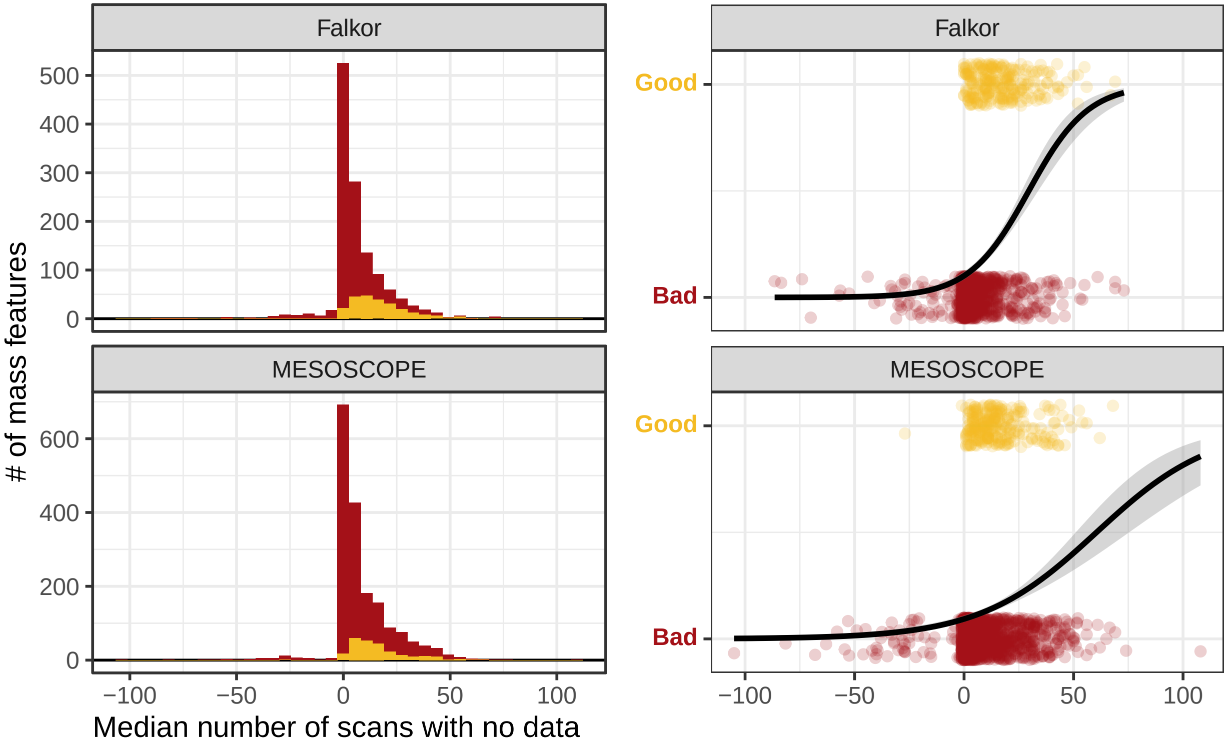
Falkor
MESOSCOPE
Falkor
MESOSCOPE
### of mass features
Median number of scans with no data
Good
Bad
Good
Bad
0
100
200
300
400
500
0
200
400
600
−100
−50
0
50
100
−100
−50
0
50
100

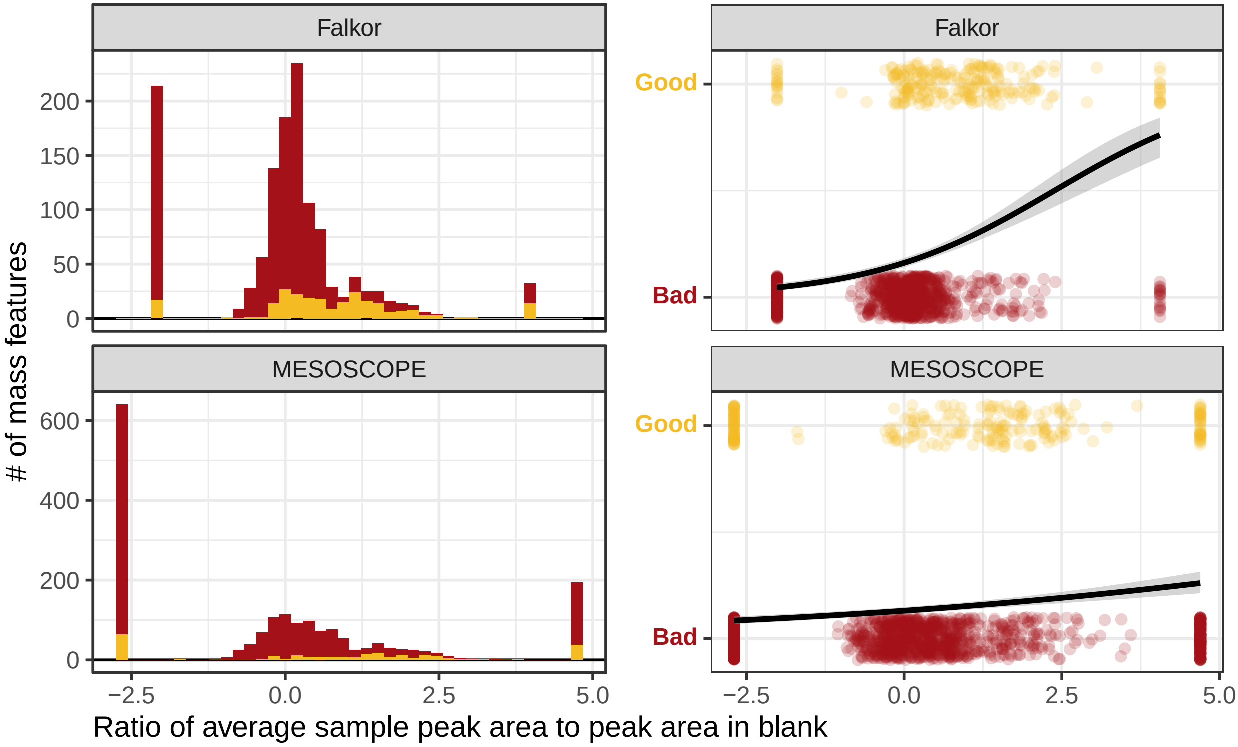

Falkor
MESOSCOPE
Falkor
MESOSCOPE
Good
Bad
Good
Bad
### of mass features
200
150
100
50
0
600
400
200
0
−2.5
0.0
2.5
5.0
−2.5
0.0
2.5
5.0
Ratio of average sample peak area to peak area in blank

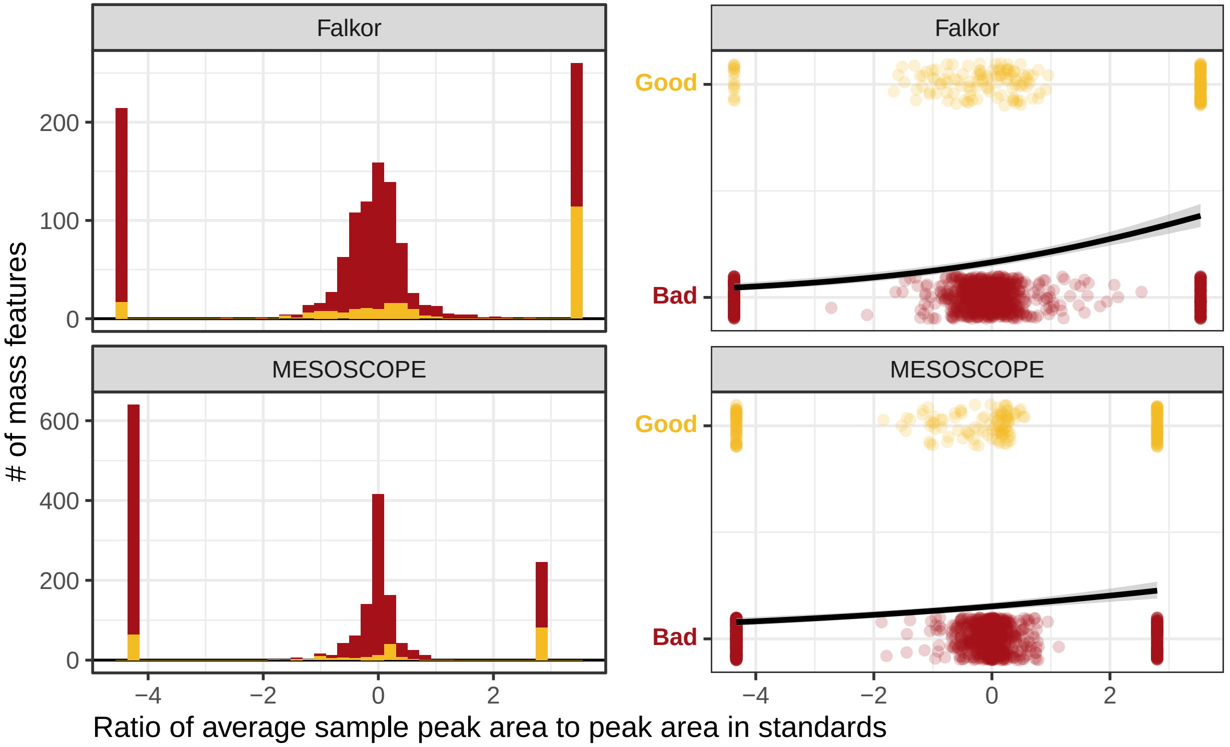

Falkor
MESOSCOPE
Falkor
MESOSCOPE
Good
Bad
Good
Bad
200
100
0
600
400
200
0
−4
−2
0
2
−4
−2
0
2
### of mass features
Ratio of average sample peak area to peak area in standards

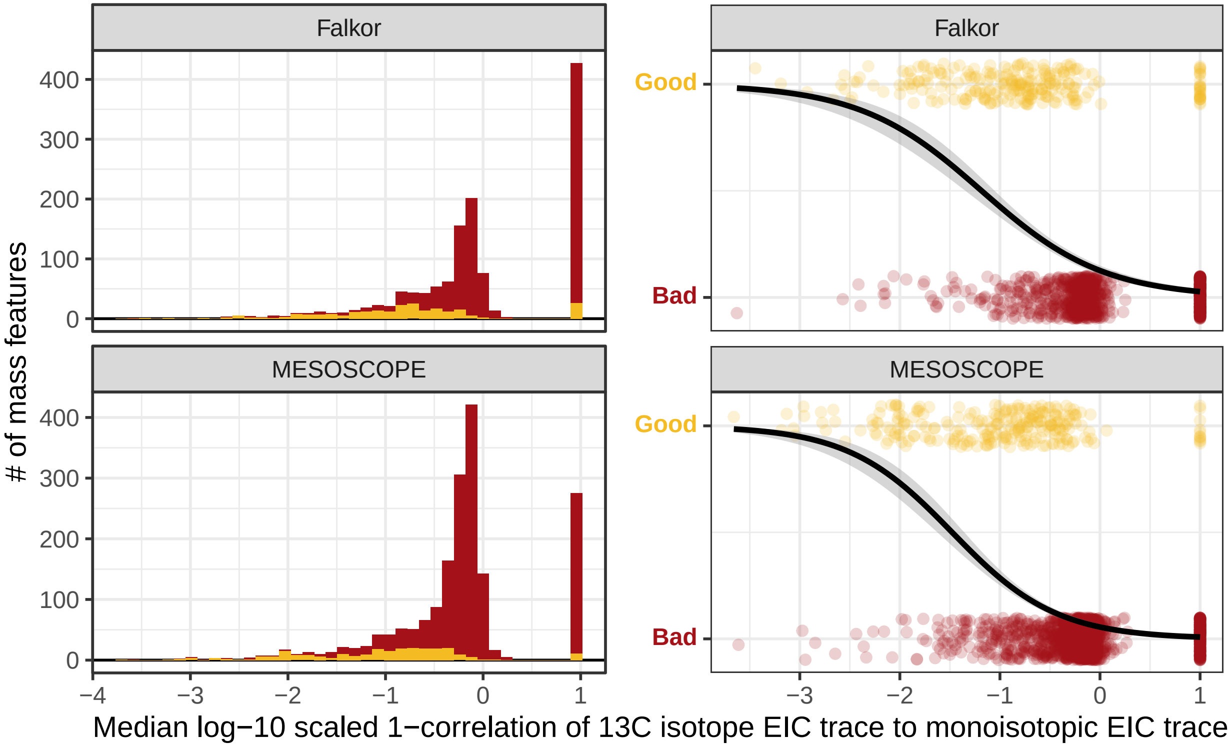
Falkor
MESOSCOPE
### of mass features
400
300
200
100
0
−4
−3
−2
−1
0
1
Good
Bad
Median log−10 scaled 1−correlation of 13C isotope EIC trace to monoisotopic EIC trace

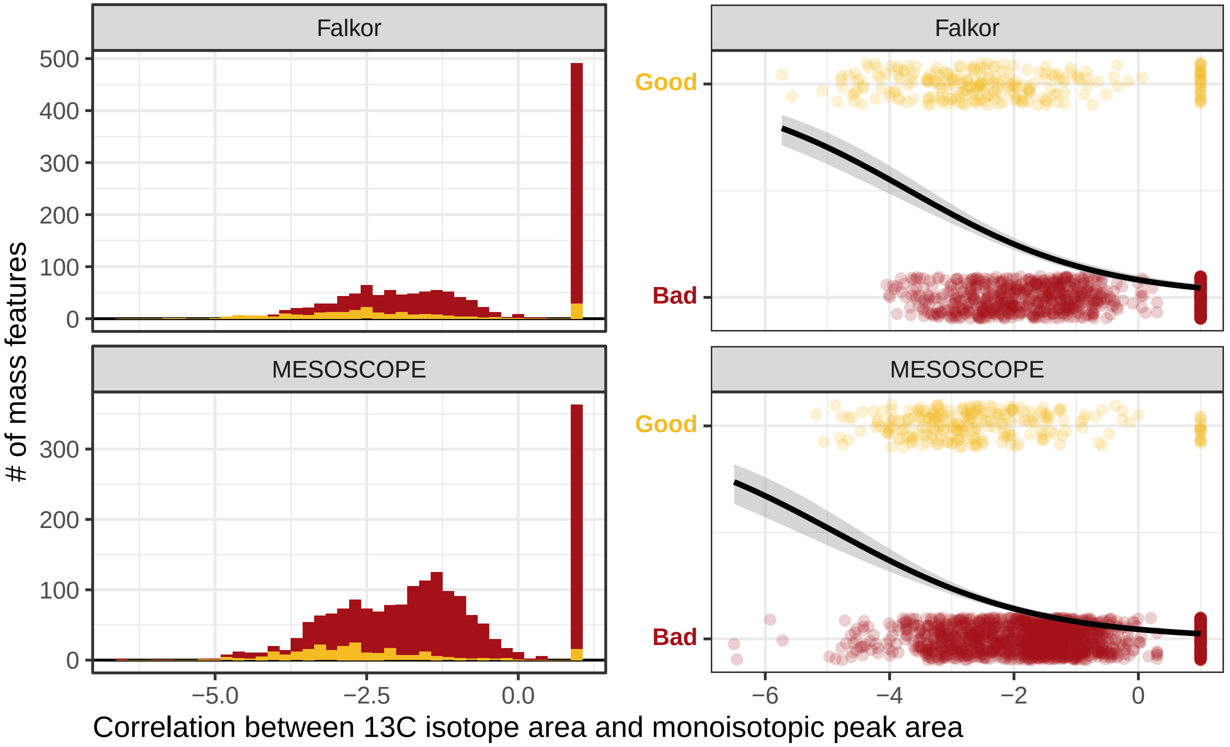

Falkor
MESOSCOPE
Falkor
MESOSCOPE
### of mass features
500
400
300
200
100
0
300
200
100
0
−5.0
−2.5
0.0
Good
Bad
Good
Bad
−6
−4
−2
0
Correlation between 13C isotope area and monoisotopic peak area
