## Supplementary figures and images for "Picky with peakpicking: assessing chromatographic peak quality with simple metrics in metabolomics"

### Supplemental Figure 2

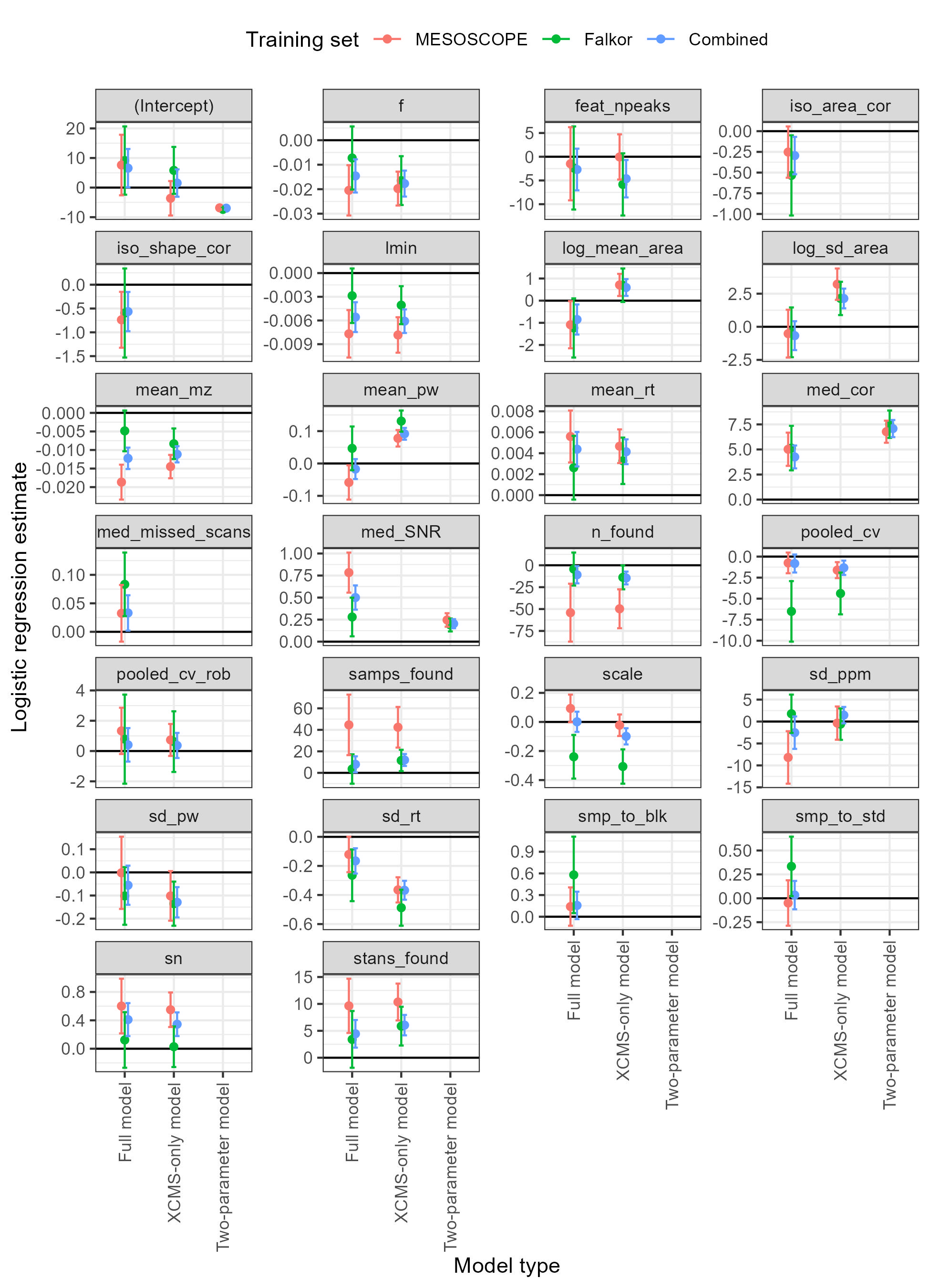

### Supplemental Figure 3

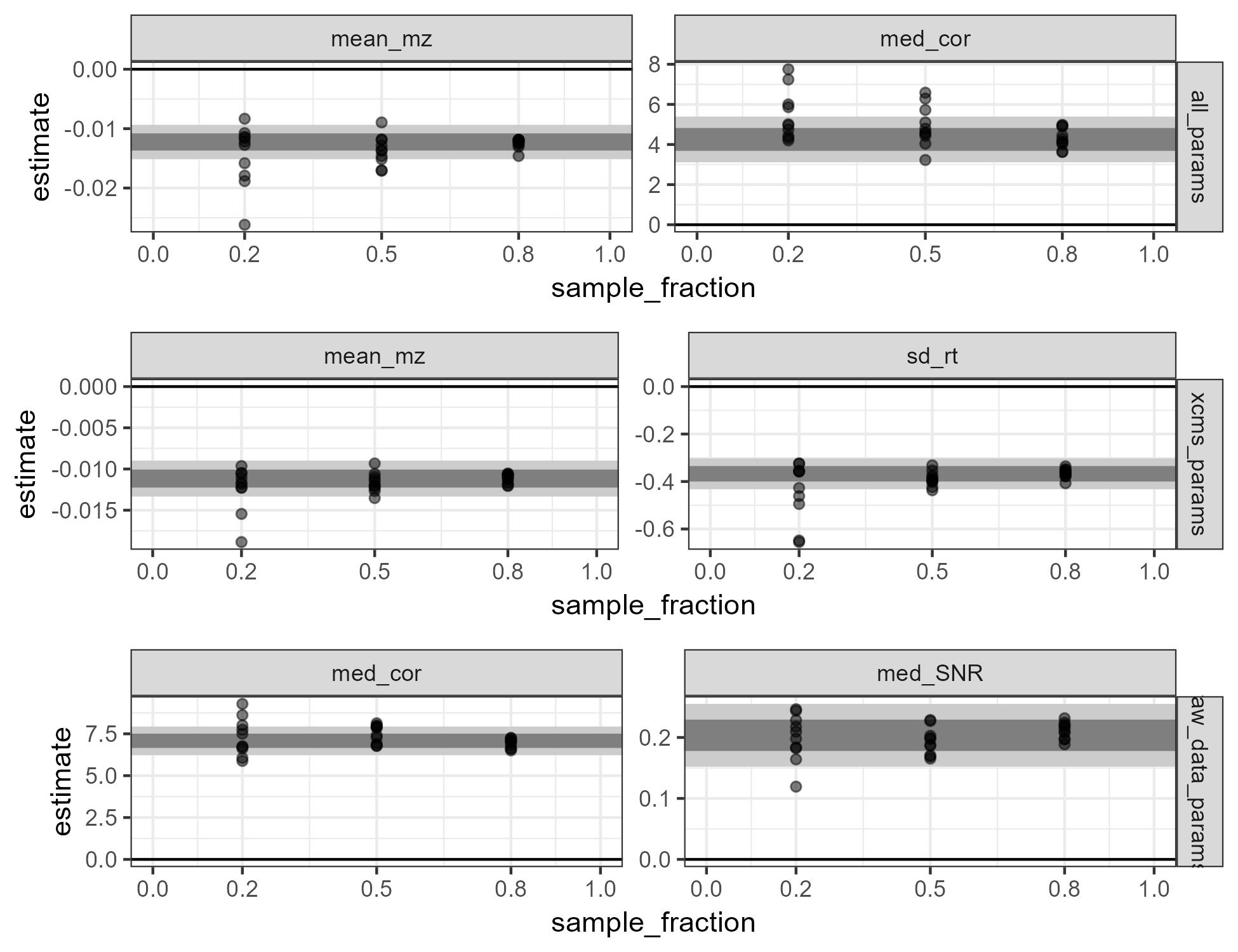

### Supplemental Figure 4

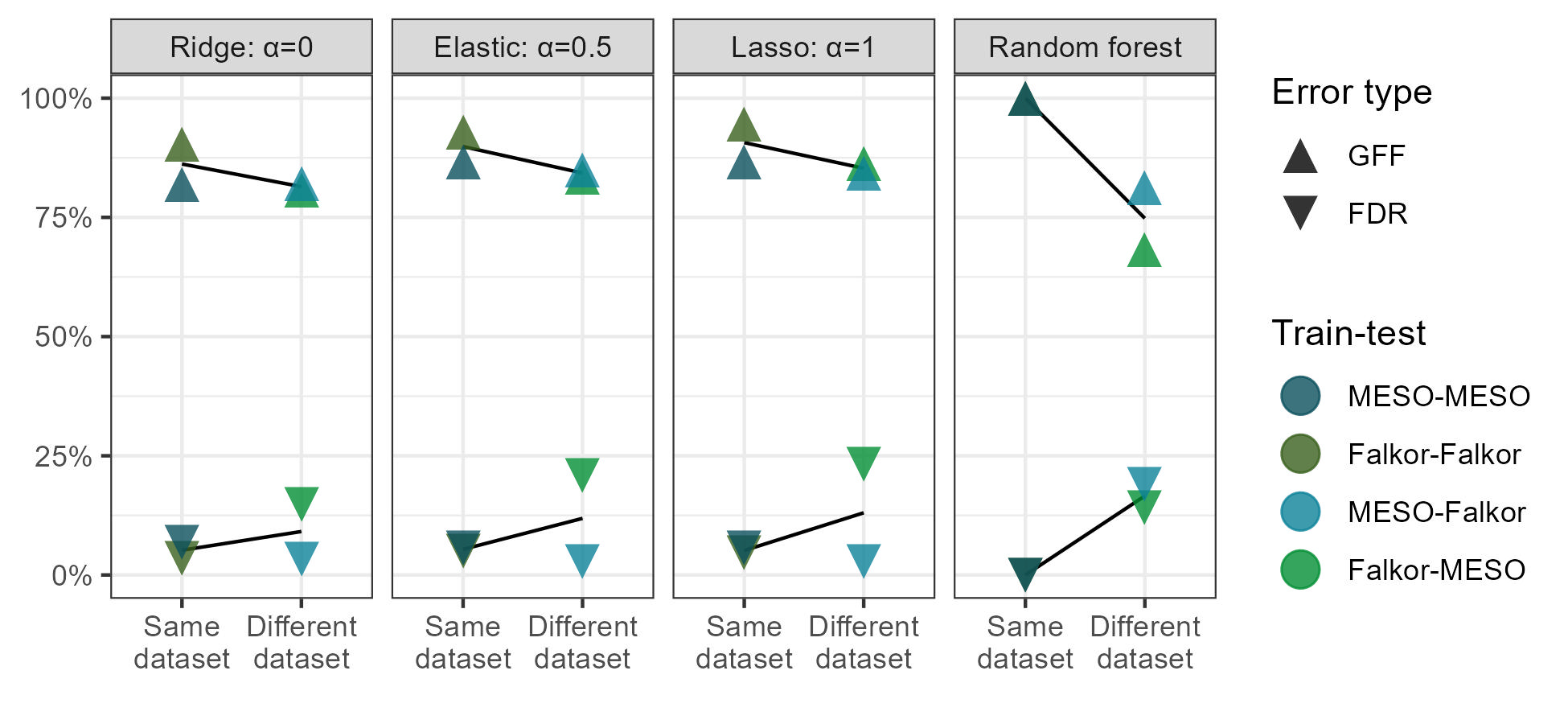
